# Physics of sliding on water predicts morphological and behavioral allometry across a wide range of body sizes in water striders (Gerridae)

**DOI:** 10.1101/2022.10.28.514156

**Authors:** Woojoo Kim, Jae Hong Lee, Thai H. Pham, Anh D. Tran, Jungmoon Ha, Sang Yun Bang, Piotr G. Jablonski, Ho-Young Kim, Sang-im Lee

## Abstract

Laws of physics shape morphological and behavioral adaptations to locomotion at different body sizes. Water striders serve as a model taxon to study how simple physical constraints of water-surface habitats affect their behavior and morphology, and hydrodynamics of rowing by midlegs on the surface is well understood. However, the physics of the subsequent passive sliding has been less explored. We created a model of sliding on the water surface to simulate the effect of body mass, striding type, and wetted leg lengths on an insect’s ability to float on the surface and on the sliding resistance. The model predicts that to support their weight on the surface during sliding, the heavy species should either develop long forelegs that support the frontal part of its body during symmetrical striding (when two midlegs thrust) or use asymmetrical striding (when one forward-extended midleg supports the body while the other midleg and contra-lateral hindleg thrust). These predictions are confirmed by the behavior and morphology of various Gerridae species. Hence, the results illustrate how simple physical processes specific to a certain habitat type have far-reaching consequences for the evolution of morphological and behavioral diversification associated with body size among biological organisms in these habitats.

## Introduction

Understanding how laws of physics may constrain morphological and behavioral evolution of biological organisms of different body sizes is of great importance not only to biology (Alexander, 1985; Garland et al., 2022; Al-Mosleh et al., 2021) but also to the modern bioinspired engineering (Yang et al., 2018; Kwak and Bae, 2018). Allometry, the study of how physics and biology affect the relationships between body size and other characteristics of an organism, has a long history (Heglund et al., 1974; Losos, 1990; Stern and Emlen, 1999; Gayon, 2000; Dial et al., 2008; Frankino et al., 2005; Labonte et al., 2016; Dodds et al., 2001; Pélabon et al., 2014; West et al., 1997; Huxley, 1932; Thompson and Thompson, 1942; Mencuccini, 2002; Smith, 1993).

Distinguishing between specific biological and physical mechanisms/constraints responsible for allometry may often be challenging (Garland et al., 2022; Dial et al., 2008). However, some organisms may provide the more clear-cut situations where allometry can be attributed to physical constraints. Animals that live on the water surface are exposed to a very clear and specific physical constraints from the nature of the water surface, and it has been suggested that body size may shape the morphological and behavioral adaptations to semiaquatic locomotion in animals (Glasheen and Mcmahon, 1996; Suter and Gruenwald, 2000; Hu et al., 2003; Hu and Bush, 2010). Water striders, Gerridae, are ideal subjects to study those issues. However, although many studies have taken theoretical approach to understand the physics of water striders’ locomotion (Hu et al., 2003; Hu and Bush, 2010; Goodwyn et al., 2008; Perez Goodwyn et al., 2009; Steinmann et al., 2018; Steinmann et al., 2021; Koh et al., 2015; Yang et al., 2016; Baek et al., 2020; Kim et al., 2017; Ma et al., 2020; Andersen, 1976; Caponigro and Eriksen, 1976; Bowdan, 1978a, b; Bühler, 2007; Lu et al., 2018; Wang et al., 2010; Bush and Hu, 2006), the research effort is confined to several small- and medium-sized water striders in spite of a wide range of body mass that spans over two orders of magnitude from less than 5 (Mahadik et al., 2020) to about 500 mg (Tseng and Rowe, 1999).

The typical locomotion mode (gait) of Gerridae comprises the ancestral symmetrical striding/skating (Perez Goodwyn et al., 2009; Andersen, 1982; Armisén et al., 2015), in which midlegs symmetrically push backwards (thrust phase) to create forward movement of the water strider body (passive sliding on the surface or leaping above the surface) while body is supported on the water surface by two forelegs and two hindlegs for the duration of the push and the subsequent sliding until the midlegs return to their original positions on the water surface and braking occurs (short-lasting braking phase). As the anterior body section remains supported on the forelegs only, the heavier the body the stronger the surface-tension force from the forelegs, otherwise the surface will break under forelegs. Hence, floating on the surface during sliding is the first theoretical consideration in predicting locomotive adaptations in large-bodied water striders. An additional consideration is the effect of body mass, wetted leg lengths (following the convention in the literature, we use the term ‘wetted length’ as water-contact length even though the leg is not technically ‘wet’ by its hydrophobicity), and sliding velocity on the resistance that the legs experience on the water surface according to general physics for water sliders (Raphaël and De Gennes, 1996; Pucci et al., 2019). Resistance may affect the efficiency of the thrust force in producing the movement and the ability of water striders to slide over long distance and duration.

Entomological literature suggests that heavy water striders evolved unique foreleg morphology and/or striding behavior in order to support the anterior part of the body on the water. Firstly, disproportionately elongated wetted forelegs in the large-bodied water striders of the genus *Ptilomera* (Andersen, 1982; Kim et al., 2022) may help to support the anterior body during the thrust and passive sliding phases. However, as this type of morphology is also observed in small species of Gerridae (e.g., in Halobatinae (Andersen, 1976; Mahadik et al., 2020), it may not necessarily be a specific adaptation to heavy body, but rather to the midlegs not being used for support on the water surface. Secondly, asymmetric striding that involves one midleg extended forward to support the heavy anterior body part while the other midleg provides thrust may be the specific adaptation to heavy body. This locomotive behavior was only reported in the world’s largest water strider species, the giant water strider, *Gigantometra gigas* (Tseng and Rowe, 1999). In the asymmetric striding, forelegs are not crucial for the body support, and *G. gigas* has relatively short forelegs. Hence, based on the above reasoning, and based on the brief review of morphological measurements of Gerridae from the literature (Fig S1), we introduce the concept of the “wetted leg geometry”. The term refers to the proportions of wetted forelegs, wetted midlegs and wetted hindlegs in the total length of the wetted legs (sum of wetted lengths of forelegs, midlegs, and hindlegs). Literature suggests that we can classify species into at least three types of “wetted leg geometry”: the “intermediate-foreleg (or “standard”) geometry” observed in the frequently studied small and mid-size genera *Gerris* and *Aquarius*, the “long-foreleg geometry” (e.g., in Halobatinae, Ptilomerinae) and the “short-foreleg geometry” (extremely developed in *Gigantometra gigas*), depending the proportion of wetted forelegs in the total length of the wetted leg (Fig S1 shows ranges of values of different taxa).

The two aspects, the support for the anterior part of the body and the resistance on the legs during the sliding, should be considered in building a theoretical model to predict the feasible combinations of the “wetted leg geometry” and striding gait (symmetric or asymmetric striding mode) for a given body mass of a water strider in a specific habitat. Here, we develop a theoretical model of the hydrodynamics of a passive sliding phase in symmetric and asymmetric striding modes for the three types of the wetted legs geometry across a range of the water strider body size. We use the model to predict allometric changes in morphological and/or behavioral adaptations to locomotion on the water surface among the species of Gerridae. The predictions can be tested in the future comparative studies once accurate behavioral and morphological data are collected.

## Results

### Theoretical model of a sliding water strider

Detailed technical explanations of the mathematical model are in the Methods and the Supplementary Materials. The model assumed the leg as a cylinder with smooth surface with the length and diameter imitating legs of water striders. We consider that the water striders can stride in symmetric or asymmetric manner with the body velocity, *U*, relative to the water surface. Hence, water striders can produce thrust symmetrically, by using two midlegs, or asymmetrically, by using one midleg and one contralateral hindleg (Fig 1A), and they can either slide symmetrically on two forelegs and two hindlegs or slide asymmetrically on a midleg and two hindlegs. When a water strider is sliding on the water surface (Fig 1B, C), the normal force (the anterior, *N*_*a*_, and the posterior, *N*_*p*_, normal force) keep the water strider afloat, while the resistance on the legs interacting with water (the anterior, *R*_*a*_, and the posterior, *R*_*p*_, resistance) gradually slows down the passively sliding water strider.

**Fig 1.**
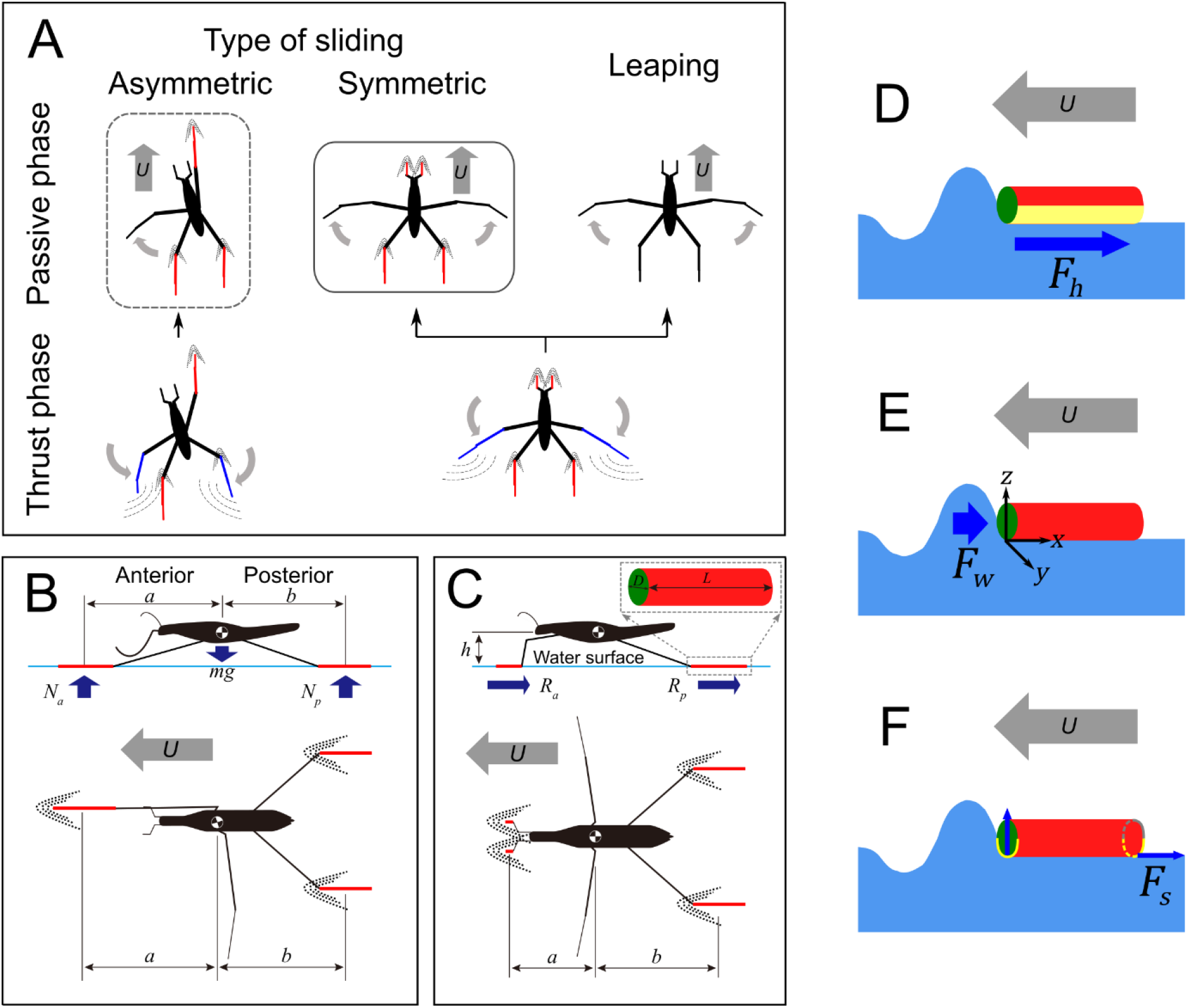
Graphical explanation of the basic concepts in the model of sliding of water striders. (A) – Striding locomotion has two phases: thrust phase (when legs pushing backward create a thrust force forward), and passive phase (when water strider is sliding on water or leaping above water). The thrust plus sliding can either be symmetric (typical for most Gerridae) or asymmetric. The leaping is preceded by symmetric thrust. Colored legs indicate thrusting legs (blue) and sliding legs (red). (B, C) – schematics of asymmetric (B) and symmetric (C) sliding, and variables used in the model: anterior and posterior normal forces (*N*_*a*_, *N*_*p*_), the anterior and posterior resistance forces (*R*_*a*_, *R*_*p*_), wetted leg lengths (*L*) and diameters (*D*), horizontal distance along line parallel to the moving direction from the center of the mass to the center of the anterior and posterior wetted legs (*a, b*), body height above water surface (*h*), body velocity (*U*); (D, E, F) – explanations of the three main forces contributing to the total resistance: hydrodynamic drag (D; *F*_*h*_), wave drag (E; *F*_*w*_), and surface tension (F; *F*_*s*_).

We assume that three types of resistance force are applied to the water strider during the passive sliding phase: hydrodynamic drag *F*_*h*_ (Fig 1D), wave drag *F*_*w*_(Fig 1E), and surface tension force *F*_*s*_(Fig 1F). We first consider the resistance force on one leg of the water strider. We assume that the leg is sliding on the water surface oriented parallel to the direction of movement (Fig 1B, C) and regardless of the water strider mass, the half of the surface of the wetted leg interacts with the water surface. The hydrodynamic drag, *F*_*h*_, is dominantly caused by the shear stress acting on the wetted area of the leg (yellow color in Fig 1D). It is a function of water properties (density, *ρ*, kinematic viscosity, *v*), leg morphology (diameter and length; the effect of the morphological structures on the surface of the leg was not considered in this study), and water strider behavior (water strider velocity, *U*, relative to the water surface). The hydrodynamic drag is greater when wetted area of the leg becomes larger and the velocity becomes faster. The capillary-gravity wave drag, *F*_*w*_, is induced by the wave on the waterfront of the cylindrical leg/water interface as shown in Fig 1E. It occurs at body velocities larger than the critical value *c* = 0.2313 m/s, when a moving water strider creates a visible wave on the water surface (also empirically proven in Fig S2). We assumed that this drag is a function of water properties (density, kinematic viscosity of water, and surface tension coefficient), morphology (body mass and leg length), and water strider behavior (body velocity).

To obtain the surface tension force that contribute to resistance (*F*_*s*_), we assumed that the slope of the water interface in front of the leg is nearly vertical (vertical blue arrow in Fig 1F) while the slope behind the leg maintains horizontal as shown by horizontal blue arrow in Fig 1F. Therefore, only the horizontal surface tension force at the posterior edge of the leg (dashed yellow half circumference in Fig 1F) contributes to the surface tension resistance, *F*_*s*_, which is a function of water property (surface tension coefficient) and leg morphology (leg diameter). We determine the resistance force on the anterior (*R*_*a*_) and posterior (*R*_*p*_) legs of a water strider as the sum of the three types of resistance (*F*_*h*_, *F*_*w*_, and *F*_*s*_).

We derived a simple gravity-normal force balance formula, and we also derived the torque-balance formula for the posterior, *N*_*p*_, and anterior, *N*_*a*_, normal forces on the legs, which depend on water strider body mass, leg morphology (distances *a, b*, and wetted leg lengths on forelegs and/or hindlegs; Fig 1A; see also Methods section), and the resistance force on the anterior and posterior legs (*R*_*a*_ and *R*_*p*_; details in the Methods). Finally, from the calculations of the system of equations from these two balance formulae, the model predicts the normal forces, the total resistance as a sum of anterior and posterior resistance on water strider legs, and deceleration caused by the resistance.

### The “wetted leg geometry” of the studied species used in theoretical calculations

Although the individuals from the six study species measured in our study followed a general allometric relationship between body mass and the total wetted leg length (Fig S4), they differed in the relative proportions of wetted foreleg (Fig 2A), midleg (Fig 2B) and hindleg (Fig 2C) lengths, and represent the three types of “wetted leg geometries”: the “intermediate-foreleg geometry”, the “long-foreleg geometry” and the “short-foreleg geometry”.

**Fig 2.**
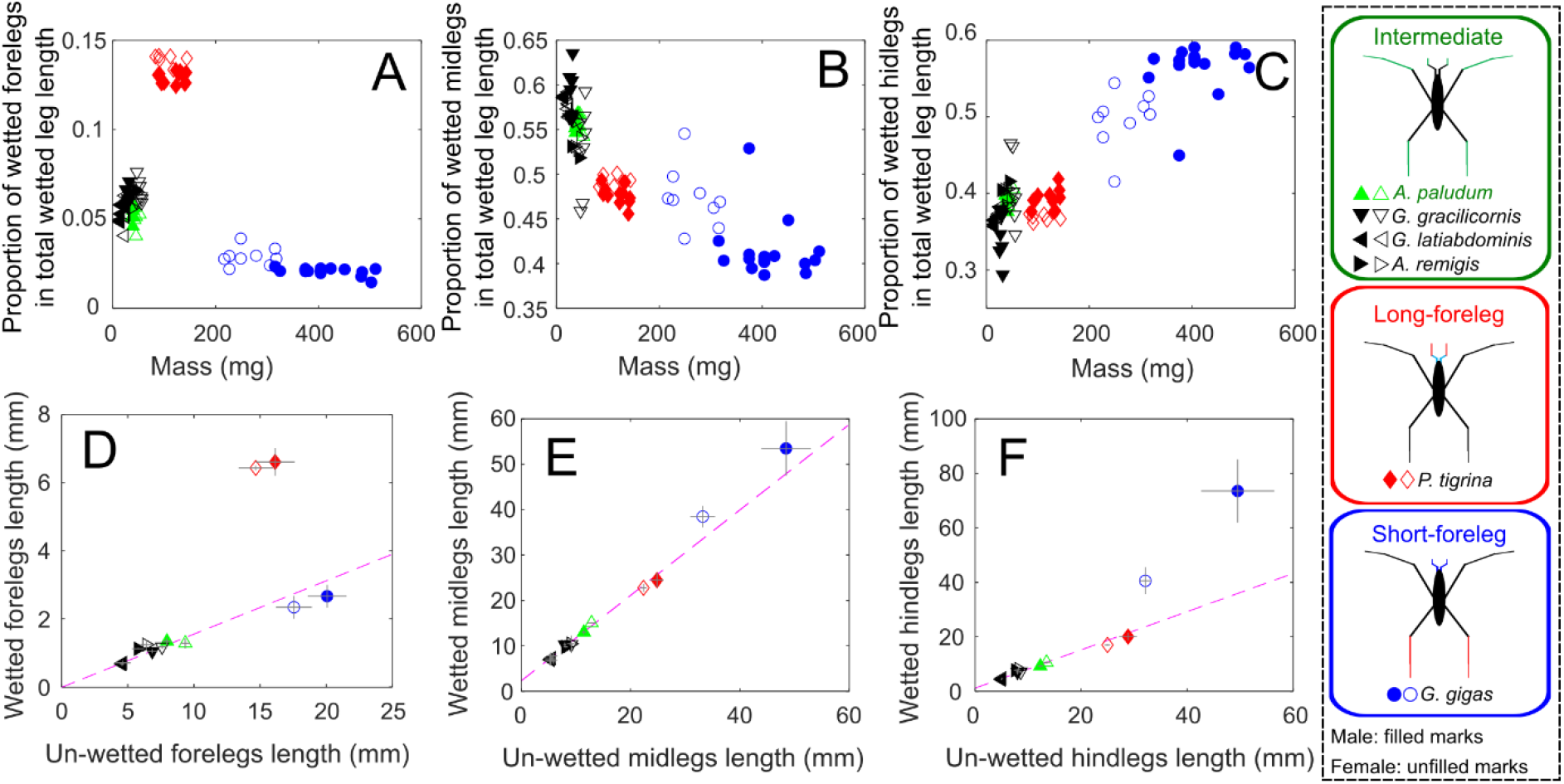
Leg proportions (“leg geometry”) and body masses of the study species. Leg morphology is expressed as proportions of wetted forelegs, midlegs, and hindlegs in the total length of wetted legs of an individual water strider (“wetted leg geometry”; A-C), and as absolute lengths of wetted *vs*. un-wetted leg for forelegs, midlegs and hindlegs of each study species (D-F). From the four species with “intermediate-foreleg wetted leg geometry” (*Aquarius paludum, A. remigis, Gerris gracilicornis and G. latiabdominis*), we chose the wetted leg geometry of *A. paludum* (green triangles) to serve as the representative distribution of leg morphology in species with “intermediate-foreleg geometry” in the model. *Ptilomera tigrina* served as the representative of “long-foreleg geometry”, and *Gigantometra gigas* served as representative of the “short-foreleg geometry” in the model.

The four small/medium size water striders that we have measured (*G. latiabdominis, G. gracilicornis, A. remigis, A. paludum*) form one cluster of “intermediate-foreleg geometry” with wetted forelegs comprising from ∼4 to ∼8% of total wetted leg length (Fig 2A-C). We decided to use the specific values of the “wetted leg geometry” of *A. paludum* (marked as green triangle in Fig 2) as the representative “intermediate-foreleg geometry” for comparisons with the two other “wetted leg geometries”: the “long-foreleg geometry” with wetted forelegs comprising 12-14% of the total wetted leg length (represented by the subtropical water striders *P. tigrina*; Fig 2A-C), and the “short-foreleg geometry” with wetted forelegs comprising 1-3% (represented by *G. gigas*; Fig 2A-C). The “leg geometries” of our study subjects are also visualized in an alternative manner in Fig 2D-F as ratios of wetted to un-wetted leg lengths.

### Model predictions for five different size classes and three leg geometries

#### General

After confirming that the theoretical model reasonably well simulates the empirically observed trajectories (Fig S3), we used it to predict how the three “leg geometries” (“intermediate-foreleg”, “long-foreleg”, and “short foreleg geometry” based directly on empirical measurements of our study species; see below “Empirical observations of the study species”) would perform in terms of floating on the water without breaking the surface during sliding, and in terms of resistance and deceleration during symmetrical and asymmetrical sliding on the water surface, in five body size classes corresponding to the recorded body mass ranges of our study species: *G. latiabdominis* (12-32 mg), *A. paludum* (35-72 mg), *P. tigrina* (83-144 mg), *G. gigas* females (217-318 mg) and *G. gigas* males (316-511 mg). This resulted in predictions for 30 situations (5 body mass classes * 2 modes of locomotion [symmetrical or asymmetrical] * 3 “leg geometries”) including 6 actually observed in our study subjects and 24 “virtual” ones that have not been recorded in our study species (Fig 3).

**Fig 3.**
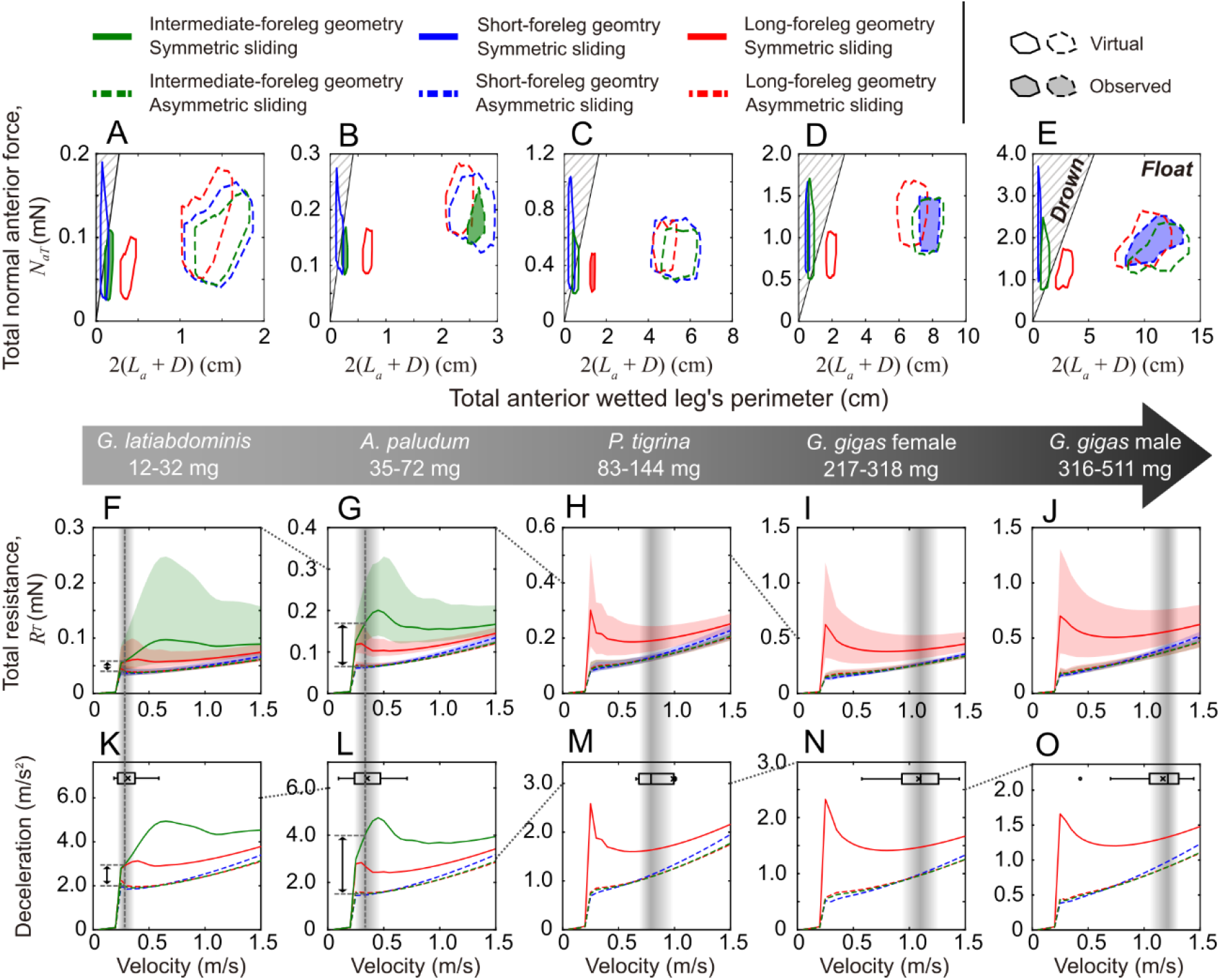
Model predictions of floating conditions, resistance, and deceleration. Model results: predictions of the ability to float on the surface without breaking it (A-E), and calculations of resistance (F-J) and deceleration (K-O) during sliding for 30 different combinations of “leg geometry” (“intermediate-foreleg”, “long-foreleg”, “short-foreleg”), body size class (5 classes between about 10 and about 500 mg) and locomotion mode (symmetric *vs*. asymmetric). The large gray arrow under the figures represents size classes based on empirical data from five species/sex classes of our study organisms: *G. latiabdominis, A. paludum, P. tigrina*, and *G. gigas* females and males. Wetted leg geometries are marked as colors: “short-foreleg” (blue), “long-foreleg” (red), and “intermediate-foreleg” (green). Sliding locomotion modes are marked with line patterns: symmetric sliding (solid line) and asymmetric sliding (dashed line). (A-E) - phase diagrams of the total normal force applied on anterior supporting legs (*N*_*aT*_ ; vertical axis) and the perimeter of the anterior legs’ wetted perimeter of the foreleg (2(*L*_*a*_ + *D*)); the diagonal black solid line in each figure corresponds to *N*_*aT*_ = 2*σ*(*L*_*a*_ + *D*), and the hatched area above the line indicates conditions leading to meniscus breaking under the anterior supporting leg(s) and sinking of the water strider’s forelegs. (F-J) The total resistance as a function of body velocity relative to the water surface for those conditions among (A-E), in which floating is possible. Geometries are marked as colors: “short-foreleg” (blue), “long-foreleg” (red), and “intermediate-foreleg” (green). Sliding types are marked with lines: symmetric sliding (solid line) and asymmetric sliding (dashed line). Empirical initial velocities observed in the study species within each size class are shown as small horizontal box plots (water striders in panels F-J), and also by vertical gray shaded rectangles across panels F-O. The resistance and deceleration differences between symmetric/asymmetric striding at the observed median velocity are marked as black arrows in F, G, K, L.

#### Theoretical predictions of conditions for floating during sliding

The maximum (critical) surface tension force that water provides to the anterior supporting leg(s) is the product of surface tension coefficient, *σ*, and the entire wetted length consisting of the length, *L*_*a*_, and diameter, *D*; (2*σ*(*L*_*a*_ + *D*)) (Koh et al., 2015; Yang et al., 2016). Therefore, the anterior supporting leg(s) would pierce through the water surface when the force needed to support the anterior part of body, *N*_*aT*_, is larger than 2*σ*(*L*_*a*_ + *D*). This force depends on multiple factors including water strider morphology, behavior, and body velocity (see details in the Methods). Using the theoretical model, we produced two-dimensional phase diagrams in Fig 3A-E, with the anterior normal force, *N*_*a*_, on the vertical axis and the wetted leg perimeter (2(*L*_*a*_ + *D*)) on the horizontal axis. In these diagrams, the conditions when the sliding water strider’s anterior supporting leg(s) do not pierce the water surface correspond to the unhatched area below the line of the critical *N*_*a*_ = 2*σ*(*L*_*a*_ + *D*). The hatched area above this line comprise situations in which the anterior supporting leg(s) will pierce the water surface.

The model predicts that the water striders do not drown if they perform asymmetric sliding in any of the 15 conditions defined by 3 leg geometries and 5 body size classes (all polygons with dashed edges in Fig 3A-E) because the wetted length of the forward-extended midleg is sufficiently long compared to the forelegs. Water striders with “long-foreleg geometry” (red solid line polygons) are not predicted to drown regardless of the body size if they perform symmetric sliding (polygons with solid line edges in Fig 3A-E). However, the three larger size classes of water striders with “intermediate-foreleg geometry” (green solid line polygons in Fig 3C-E) and water striders with “short-foreleg geometry” regardless of the body size (blue solid line polygons) are predicted to drown if they perform symmetric sliding except for a very narrow range of conditions that locate them under the critical lines (*N*_*a*_ = 2*σ*(*L*_*a*_ + *D*)).

#### Theoretically calculated sliding resistance

We calculated the relationship between body velocity and the total sliding resistance force (Fig 3F-J) for all floatable conditions. The total sliding resistance depends on the body mass, leg geometry, sliding mode, and body velocity. In general, the asymmetric sliding (broken lines in Fig 3F-J) generates lower resistance than the symmetric sliding (solid lines in Fig 3F-J). The sliding resistance dramatically increases when body velocity exceeds the minimum threshold velocity at which surface waves are produced by legs sliding on water surface (*c* = 0.2313 m/s). For symmetric sliding, the resistance greatly depends on the leg geometry and body mass: sliding resistance increase reaches a peak of 0.1-0.2 mN for body speeds of about 0.5 m/s for “intermediate-foreleg geometry” in water striders from the two smaller size classes (green solid line in Fig 3F, G) and a peak of 0.3-0.6 mN for body speed of about 0.25 m/s for “long-foreleg geometry” in water striders with large body weight (red solid line in Fig 3H-J), while for smaller water striders this peak for “long-foreleg geometry” is much less pronounced (red solid line in Fig 3F, G). For asymmetric sliding (broken lines in Fig 3F-J), a steep increase in resistance is predicted as body velocity passes through the threshold critical velocity, *c*, and afterwards its slope becomes much milder (broken lines in Fig 3F-J), but these patterns were predicted regardless of the leg geometry.

During the thrust phase of each stride, a water strider must create a total thrust force comprising a counter-resistance component (to overcome the resistance) and a net thrust force that contributes directly to the water strider body’s momentum change (and produces the body velocity observed at the start of the sliding phase). We compared the empirically estimated net thrust forces in a set of strides by our study species with the theoretically calculated resistance in those strides (Fig S5, S6), and found out that on average 85-95% of total thrust is converted into the water strider’s body momentum. The remaining thrust is used up to overcome the resistance (Fig S5, S6), especially in the symmetrical sliding of *P. tigrina* and those striders that perform fast sliding and produce surface waves (above the critical velocity threshold of 0.23 m/s) when the wave resistance starts affecting the moving water strider (Fig S5, S6).

#### Theoretically calculated sliding deceleration

As the resistance mainly contributes to the deceleration, the patterns of theoretically calculated deceleration were similar to those of the resistance. The theoretically predicted decelerations were smaller for asymmetric than for symmetric sliding for all five size classes (Fig 3K-O). The deceleration experienced by the two smaller studied species at their actual sliding velocities (shown as horizontal box-and-whiskers plots in Fig 3F, G, and marked by vertical gray shaded bars across Fig 3F, G, K, L) ranges from less than 2 to 5 m/s^2^ for symmetrical sliding and between 1.5 and 2 m/s^2^ for asymmetrical sliding (the ranges are based on lower and upper quartile values of sliding velocity recorded in the species). As the average sliding velocity of these species is less than 0.5 m/s (Fig 3F, G), these values of decelerations have relatively strong slowing-down effect compared to the larger species (see below). Additionally, the theoretically predicted difference between asymmetrical and symmetrical sliding in the deceleration at the empirically measured median body velocity is roughly twice as large in the medium-size *A. paludum* as it is in the small-size *G. latiabdominis* (black double arrows in Fig 3K, L), indicating that by using asymmetrical rather than the symmetrical striding the medium-size species with “intermediate-foreleg geometry” may importantly increase its sliding performance.

The predicted deceleration experienced by the three studied large species/sex classes at their actual sliding velocities (shown as horizontal box-and-whisker plots in Fig 3H, I, J and marked by vertical gray shaded bars across Fig 3H-J, M-O) vary ∼0.8-1 m/s^2^ in the *G. gigas* males and females to ∼1-1.5 m/s^2^ in *Ptilomera*. As the average sliding velocity of the two larger classes is more than 1 m/s (Fig 3I, J), these relatively small values of decelerations have relatively weak slowing-down effect, compared to the effect of deceleration expected in the two smaller species.

### Empirical observations of the study species

We observed three combinations of thrusting-sliding phases of locomotion: symmetric thrusting – symmetric sliding (Fig 4A, B, C), symmetric thrusting – leaping (Fig 4A, B), and asymmetric thrusting – asymmetric sliding (Fig 4B, D, E). The smallest species with “intermediate-foreleg geometry”, *G. latiabdominis*, thrusts symmetrically (except for changing direction of the body), and slides symmetrically or leaps forward after symmetric thrusting (Fig 4A; Movie S1; movie content descriptions and examples of digitized strides are in Figure S7-S10). The larger species with “intermediate-foreleg geometry”, *A. paludum*, used all three phase combinations (Fig 4B; Movie S2) depending on their initial body velocity (the velocity at the end of the thrust phase). The large species with “long-foreleg geometry”, *P. tigrina*, used only the symmetric thrust followed by symmetric passive phase (Fig 4C; Movie S3). Only when forelegs were handling the food (Kim et al., 2022) or grooming (Movie S4), *P. tigrina* used asymmetric striding mode. Both sexes of the large species with “short-foreleg geometry”, *G. gigas*, used only the asymmetric locomotion mode (Fig 4D, E; Movie S5): at least one middle leg always supported anterior part of the body even in changing direction of the body (Movie S6).

**Fig 4.**
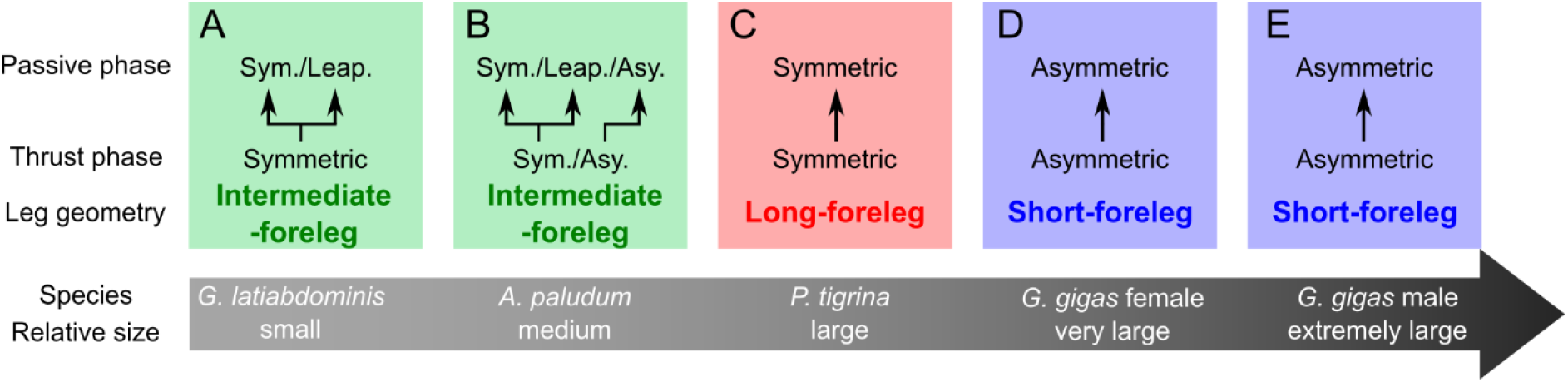
Summary of observations of locomotion of the study species (see Movies S1-S5). The large gray arrow under the figures represents relative size of species. Geometries are marked as colors: “short-foreleg” (blue), “long-foreleg” (red), and “intermediate-foreleg” (green). Water strider size classes are arranged in order from the smallest to largest size: (A) *G. latiabdominis*; (B) *A. paludum*; (C) *P. tigrina*; (D) *G. gigas* females and (E) *G. gigas* males. Supplementary movies illustrate the natural striding locomotion of *G. latiabdominis* (Movie S1), *A. paludum* (Movie S2), *P. tigrina* (Movie S3), and *G. gigas* (Movie S5). Additionally, Movie S4 illustrates that asymmetrical use of midlegs is observed in *P. tigrina* when it cannot use forelegs to stride during self-grooming behavior.

The two smaller species with “intermediate-foreleg geometry” moved at relatively slower velocities than the three larger size classes using either symmetric or asymmetric mode (horizontal plots at the top of each panel in Fig 3F-J; See also Fig S11). The largest class (asymmetrically sliding *G. gigas* males with “short-foreleg geometry”) moved with the highest speed (compare the horizontal box and whiskers plots inserted in Fig 3H, I, J; see also Fig S11). The medium size species, *A. paludum*, was observed to slide at the widest range of body velocities from near zero to near 1.5 m/s (horizontal box-and-whisker plot in Fig 3G and Fig S11).

### Locomotion mode depends on body speed – observations in *G. latiabdominis* and *A. paludum*

Observations of *G. latiabdominis* revealed that they used symmetric thrust followed by either sliding or leaping (Fig 4A). Leaping velocity of *G. latiabdominis* was significantly faster than that of symmetric sliding (Wilcoxon Signed-Rank Test, p<0.05, n=8, Table S2). Hence, they seemed to avoid sliding on the water surface by leaping in conditions of high resistance, i.e., when they perform forward locomotion at high body velocity.

Observations of *A. paludum* revealed that symmetric thrust followed by leaping occurred mostly at high body velocities (>0.5 m/s for *A. paludum*; Fig 5A, S12). *A. paludum* switched between symmetric and asymmetric modes of sliding locomotion (Fig 4B), i.e., the same individual sometimes used the symmetric and sometimes the asymmetric striding. Asymmetric mode was used during forward locomotion that at the final moment of the thrusting phase was statistically significantly faster than during symmetric striding mode (Fig 5A, S12; statistics in Table S3). While these differences in the body velocity at the initial stage of sliding are statistically significant, symmetric sliding was used over a relatively wide range of body velocities including slow sliding (Fig 5A, S12). In contrast with the symmetric sliding, most of the asymmetric sliding occurred at the body velocities that were larger than the theoretical threshold velocity (*c* = 0.231 m/s; marked with red unfilled circle in Fig 5A, D), above which capillary-gravity wave resistance starts to slow down the water striders, especially during the symmetric sliding (compare resistance and deceleration during symmetric and asymmetric sliding in Fig. 3G, L, 5D). 75% of asymmetric sliding occurred at the initial velocities higher than 0.258 m/s (lower quartile in Fig 5A; marked by the red arrow in Fig 5A, D) when symmetrical sliding already results in twice as strong deceleration due to resistance as the asymmetrical sliding does (Fig 5D, red double arrow shows this difference). Higher body velocity leads to an increasingly larger difference in resistance between symmetric and asymmetric striding (Fig 5D). It is illustrated by the relatively smaller predicted deceleration difference between symmetric and asymmetric sliding for the velocity ∼0.37 m/s, corresponding to the median initial velocity of symmetric sliding (marked with the green filled triangle on the velocity axes in Fig 5A, D), and the relatively larger deceleration difference for the higher body velocity of ∼0.44 m/s corresponding to the median (marked with the un-filled green triangle on the velocity axes in Fig 5A, D) initial velocity of asymmetric sliding (these differences in deceleration are marked with green solid and green broken arrows in Fig 5D). The lower deceleration in asymmetric sliding is associated with the sliding distance (Fig 5B; Table S4) and sliding duration (Fig 5C; Table S5) twice as long for asymmetric as for the symmetric sliding (detailed results of all the statistical analyses are in Tables S3-S5). However, this seems not to be a pure mechanical outcome of lower resistance because the end of sliding (distance, duration) is largely dependent on the moment at which the water strider decides to put down its midlegs on the water surface, which practically defines the end of sliding as seen in examples of locomotion shown in Movie S2.

**Fig 5.**
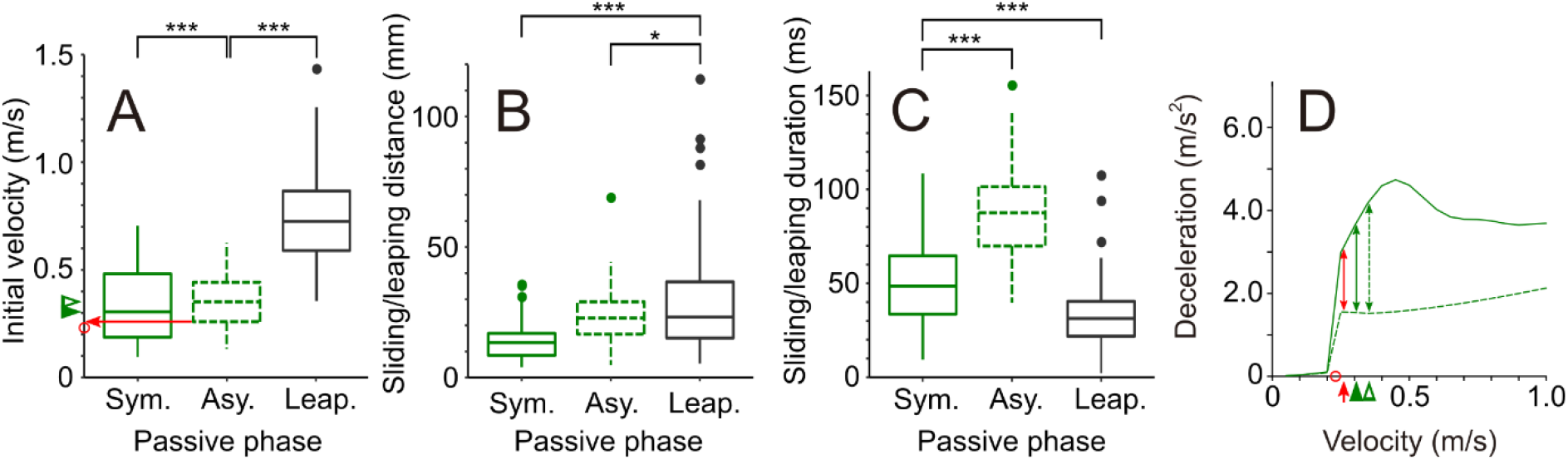
Striding behavior of *A. paludum* and theoretical predictions of deceleration for *A. paludum*. The box plot of initial velocity (A), sliding/leaping distance (B), and sliding/leaping duration (C) in a passive phase from empirical data of *A. paludum*. Symmetric sliding, asymmetric sliding, and leaping are marked as green solid, green dashed, and gray solid lines, respectively. Theoretical predictions of deceleration in (D) come from Fig 3L. Median initial velocities of symmetric and asymmetric sliding are marked with filled and unfilled triangles in (A, D), respectively. Lower quartile of asymmetric sliding velocity is marked with red arrow in (A, D). Critical body velocity of wave-making, *c* = 0.231, is marked with unfilled red circle in (A, D). The statistical analysis results for (A-C) are in Tables S3-5, and additional results for *A. paludum* are in Fig S13 and Table S6.

## Discussion

Our analysis predicts that all six combinations of the three leg geometries and the two locomotion modes can theoretically be observed among the relatively small-sized water striders (∼10 to ∼30 mg; considering floating ability during sliding). Although the symmetrical locomotion by the water striders with “short-foreleg geometry” is physically possible (i.e., water striders can stay afloat), this can be performed only in a narrowly constrained area of light body mass or slow motion (i.e., when a relatively weak normal force is applied on the anterior body part) in these small-sized water striders, because if the body is heavier and/or locomotion is faster, the short wetted forelegs cannot create sufficiently large force upward to support the anterior body section on the water surface during sliding. This also indicates that very small water striders (smaller than the body size modeled here, i.e., <10 mg) could theoretically perform symmetrical mode of locomotion with the “short-foreleg geometry”.

However, as those theoretically feasible combinations differ in resistance, and the resulting deceleration, and as water striders seem to pay attention to the resistance (as indicated by our observations of the behavioral plasticity in *A. paludum*), we hypothesize that natural selection or adaptive behavioral plasticity towards decreasing resistance may in certain conditions cause evolutionary or behavioral shifts from the ancestral (Perez Goodwyn et al., 2009; Andersen, 1982; Armisén et al., 2015) symmetric striding of water striders with “intermediate-foreleg geometry” towards either the asymmetric locomotion mode or “long-foreleg geometry”. Asymmetric locomotion mode substantially decreases resistance and deceleration and increases sliding distance but involves weaker thrust from only one midleg aided by contralateral hindleg. Hence, it is feasible only in habitats where the relatively strong thrust and frequent striding are less important (stagnant or slow-flowing water). The “long-foreleg geometry” reasonably decreases resistance (longer forelegs create lower resistance by smaller wave drag) while maintaining high thrust from two symmetrically pushing midlegs, which will be especially important in ecological situations where frequent rowing with high thrust is highly beneficial (e.g., in fast flowing water). The presence of “long-foreleg geometry” (and apparently also the symmetrical gait) even in the small taxa typical for fast current (e.g., *Metrocoris*; Perez Goodwyn et al., 2009) or for turbulent oceanic waters (*Halobates*; Andersen, 1976; Mahadik et al., 2020) is consistent with the idea that “long-foreleg geometry” is advantageous in the turbulent habitat where frequent thrust from midlegs is needed even in the smaller water striders.

When body mass reaches the range represented by *P. tigrina* and *G. gigas* (range of about 80-500 mg), water striders with typical “intermediate-foreleg geometry” of legs would not be able to support their bodies on the surface during symmetric striding/sliding (when the anterior body mass is supported by two forelegs). The model predicts, and literature (Matsuda, 1960) suggests (Fig 6 and Fig S14), that there are two solutions. One solution involves a shift to “long-foreleg geometry” by elongation of forelegs (recent studies in the genetics of morphology in Gerridae identified some genes that may be involved in the leg elongation; Khila et al., 2009; Refki and Khila, 2015) while maintaining the standard symmetric locomotion mode like in Ptilomerinae. The other solution involves the use of asymmetric locomotion mode, like in *G. gigas*. The difference between *G. gigas*, who lives in slower flowing waters, and *P. tigrina*, who lives in faster flowing water, is consistent with the idea that even though the asymmetric sliding always creates less resistance than the symmetric sliding and does not cause sinking regardless of body mass and leg geometry, *P. tigrina* does not use the asymmetric sliding (exception shown in Movie S4) because of the importance of strong thrust in the very frequent short strides against the fast flowing water in their habitat (Kim et al., 2022; see also Movie S3). Hence, we propose that the habitat type may affect the evolutionary trajectories shaping the wetted leg geometry in large water striders leading to the asymmetrical locomotion in slow-flowing waters or to the long-foreleg/symmetrical locomotion combination in species from fast currents, where the requirements for frequent and strong thrust may additionally trigger evolution of special micro-structures for rowing (Mahadik et al., 2020; Kim et al., 2022; Polhemus and Zettel, 1997) and the associated loss of the midlegs’ function of supporting the water strider on water surface (Andersen, 1976; Kim et al., 2022). If this is correct, then Gerridae illustrate how the physical environment channels the morphological and behavioral evolution (Dobzhansky, 1974; Martínez and Moya, 2011) towards either of the two physically feasible adaptive solutions for locomotion by large-sized water striders.

**Fig 6.**
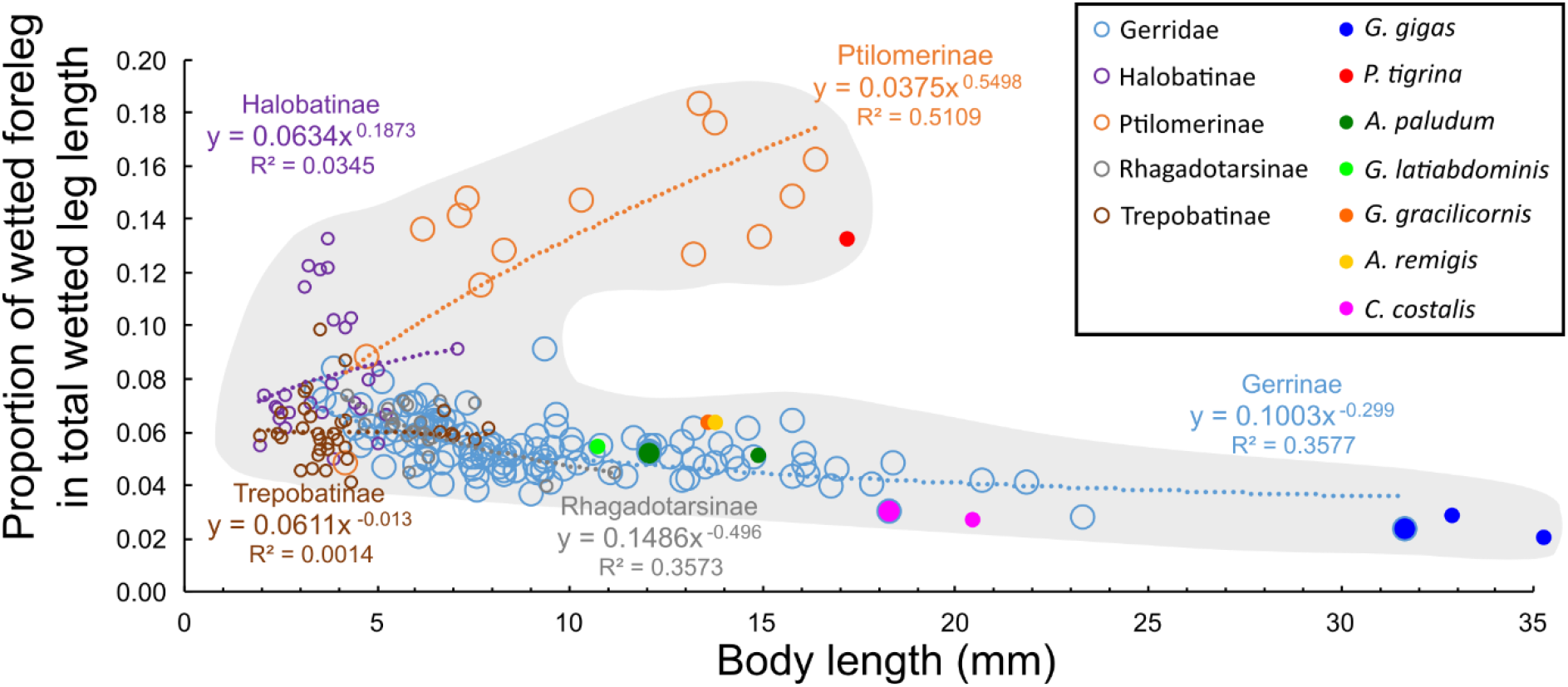
The body length and the proportion of wetted foreleg length in total wetted legs’ length. The gray shaded area helps visualizing that with increasing body size the water striders adopt one of the two “wetted leg geometries”, either “long-foreleg” or “short-foreleg geometry”. The subfamilies (Matsuda 1960) are indicated by different large unfilled circles: Gerrinae (blue), Ptilomerinae (orange), Halobatinae (purple), Rhagadotarsinae (gray), and Trepobatinae (brown). The species with our measured data are indicated by small-filled circles: *G. gigas* (blue), *P. tigrina* (red), *A. paludum* (green), *G. latiabdominis* (light green), *G. gracilicornis* (orange), *A. remigis* (yellow), and *C. costalis* (purple). The subfamilies follow Matsuda 1960, which may be not entirely consistent with the modern assignments of genera into subfamilies. Additionally, this is phylogenetically un-corrected relationship, and therefore it does not directly represent evolutionary processes shaping the evolutionary changes of leg morphology as a function of evolutionary changes of body size (the goal of the future studies). The figure is based on Table 16 in Matsuda 1960. See also Fig S14 for more detailed comments about Table 16 in Matsuda (1960). The equations fitted to the data points for each family separately have the power form following the convention for allometric equations. However, we used body length because of the absence of data for body mass, absence of body width, height or diameter data, and absence of body length – body mass formulas for water striders over such a large body size range. We expect that from among the possible linear measurements of body (width, height, length) the length is relatively more correlated with the body mass (albeit not necessarily in a linear fashion) than are body width or height as they are relatively small and differ among species relatively less than the body length. We decided not to use *body length*^*3*^ (a possible alternative used occasionally in allometry) because of the elongated shape of the water striders.

Asymmetrical mode provides a similar performance regardless of the relative wetted foreleg length and therefore it is not surprising that, in accordance with the rules of competition among developing body parts (Nijhout and Emlen, 1998), it is associated with shortening of the wetted forelegs that are no longer needed for support of the anterior body mass like in *Gigantometra gigas* and most likely in other large Gerrinae with “short-foreleg geometry” (Fig 6 and Fig S14). Finally, as already speculated (Tseng and Rowe, 1999), the asymmetric locomotion is associated with asymmetry in thrust (stronger on the side of the pushing midleg than on the side of the midleg stretched forward), which leads to torque in the horizontal plane. Therefore, the especially elongated wetted hindlegs characterizing the “short-foreleg geometry” (Fig S14C) of the asymmetrically striding species may play a role as a rudder preventing rotation of body axis. If this is correct, the hindlegs in heavy asymmetrically striding species serve two functions: adding to the thrust and counteracting the torque.

As we have discovered asymmetric locomotion mode in one of the common and widespread species, *A. paludum* (Andersen, 1982), which has been the subject of multiple studies (Goodwyn et al., 2008; Koh et al., 2015; Yang et al., 2016; (Crumière et al., 2016; Goodwyn and Fujisaki, 2007), and was believed to solely use the symmetric locomotion mode, we advise caution in using the traditional knowledge (in the literature) about locomotion modes of water striders in natural situations. Additionally, the data on the species-specific body size usually includes information on body length but not fresh body mass, and the existing literature on body length-body mass relationships in insects does not concern fresh body mass (Smock, 1980; Ganihar, 1997; Poepperl, 1998; Analytics and Cressa, 1999; Grandgirard et al., 2002; Gilbert, 2011; Martin et al., 2014; Nakagawa and Takemon, 2014; García -Barros, 2015) or it does not provide accurate formulas for the body shape and the full body mass range of Gerridae (Sage, 1982; Brady and Noske, 2006). Hence, the information from the literature allows us to present only a very preliminary view on the relationship between relative body length and “wetted leg geometry” (Fig 6 and Fig S14), which nevertheless confirms the model predictions. Our preliminary observations of a relatively little studied genus of large water striders, *Cylindrosthetus costalis* with “short-foreleg geometry” (Fig 6), confirms that, similar to *G. gigas*, they use asymmetric striding mode (Movie S7). Once solid morphological and behavioral data on locomotion modes in natural habitats across a variety of species of different sizes are collected, the predictions from our theoretical model can be properly tested in quantitative comparative phylogeny-based analyses of evolutionary correlations between body size, morphological adaptations (leg geometry) and behavioral plasticity (locomotion mode), in a variety of habitats across a wide range of body weights from less than 5 mg (e.g., Halobatinae) to above 500 mg in *G. gigas*. Hence, the model provides a solid theoretical basis for the next comparative step of research to understand the evolution of allometry of striding in water striders. It also provides insights into bio-inspired engineering of water walking robots of various sizes (Kwak and Bae, 2018; Koh et al., 2015; Wang et al., 2010; Hu et al., 2007; Shin et al., 2008).

## Materials and Methods

### Mathematical Model

The total resistance of each leg was calculated based on three forces: hydrodynamic drag (White, 2010; Fig 1D), wave drag (Fig 1E), and surface tension (Fig 1F). The normal forces on a leg (Fig 1B) supporting the anterior side (*N*_*a*_) and the posterior side (*N*_*p*_) were calculated by the force balance in the vertical direction and the torque balance about the center of the mass of the water strider. We computationally determined *N*_*a*_, *N*_*p*_, *R*_*a*_, and *R*_*p*_ (Fig 1B, C) for an empirical situation of an water strider sliding on the surface comprising the following set of empirically derived values: body mass, *m*, wetted leg lengths, *L*, wetted leg diameters, *D*, distances *a* and *b* (as defined in Fig 1B, C), vertical distance between surface and water strider body, *h*, (Fig 1C) and body velocity, *U*, during sliding.

The larger the resistance force is, the larger the rate of deceleration is pronounced, and the heavier the water strider is, the smaller the rate of deceleration is. We have estimated the deceleration rates (Fig 3K-O) corresponding the lines of average resistance in all five panels in Fig 3F-J. We calculated the body deceleration for the average mass of each size class from the standard equation: 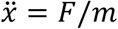 with *F* being the total external force.

#### Hydrodynamic drag (*F*_*h*_; Fig 1D)

The hydrodynamic drag on a cylindrical leg is a function of water properties (density and kinematic viscosity of water), morphology (diameter and length of the wetted leg), and insect behavior (water strider velocity, *U*, relative to the water surface). The hydrodynamic drag, *F*_*h*_, is dominantly caused by the shear stress on the leg surface in contact with water (yellow color in Fig 1D), and is represented as

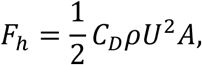

where *ρ* is the density of water, *U* is the relative velocity of the water strider to the water, and *A* is the wetted area of the leg (the yellow-shaded part of the cylindrical wetted leg in Fig 1D). The wetted area is assumed as a half of the curved surface of a cylinder, *A* = *πDL*/2. *D* and *L* are the diameter and the length of the wetted leg, respectively. *C*_*D*_ is the drag given by 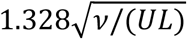, where *v* is kinematic viscosity of water (White, 2010). The resistance by hydrodynamic drag is higher when wetted area of the leg becomes larger and the velocity becomes faster.

#### Wave drag (*F*_*w*_; Fig 1E)

The capillary-gravity wave drag on a cylindrical leg is a function of water properties (density, kinematic viscosity of water, and surface tension coefficient), morphology (body mass, leg length and shape of the wetted area), and water strider behavior (water strider velocity, direction of movement relative to the leg orientation). When a floating object moves on the surface of water at velocity greater than *c* = (4*gσ*/*τ*)^1/4^, where *σ* is the surface tension coefficient of water and *g* is the gravitational acceleration, it generates capillary-gravity waves [45]. The theoretical minimum critical velocity, *c*, that produces those waves on water is 0.2313 m/s, and observations of water striders are generally consistent with this value of *c* (Fig S2).

The wave drag, *F*_*w*_, is induced by the wave generated by the cylindrical leg as shown in Figure 1e. The waves push the leg of the water strider with a force of

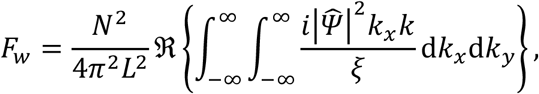

where *N* is the normal force on the leg from the water (Fig 1B) and *k*, the wave number, is represented for each x and y axis in Fig 1E as 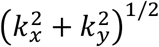 (*ℜ* stands for “real part of”). *ξ* is represented as below to simplify the formula.

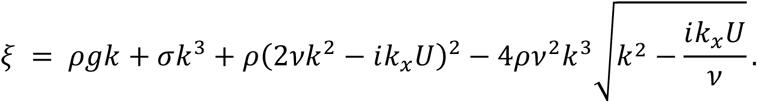

The shape of wetted area, *Ψ*, depends on the shape of the leg and its moving direction (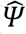 stands for the Fourier transform of *Ψ*). We assume the shape of the wetted leg as a line with length *L* and the longitudinal movement, then 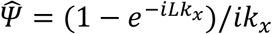. The resistance by wave drag is higher when the normal force is higher and the leg length is shorter.

#### Surface tension resistance force (*F*_*s*_; Fig 1F)

The surface tension force contributing to the total resistance during sliding is a function of water properties (surface tension coefficient) and morphology (leg diameter). For simplicity, our model assumes that the resistance by surface tension is a stepwise function that has zero value below the minimum velocity at which surface waves are produced, *c* = 0.2313 *m*/*s*, and increases beyond this threshold. As the waves are generated around a leg, we assumed that the slope of the water interface in front of the leg approaches the vertical while the slope behind the leg maintains horizontal as shown by blue arrows in Fig 1F. Therefore, only the horizontal surface tension force at the posterior end of the leg, *F*_*s*_, contributes to the resistance force. We assumed that the wetted length for the horizontal *F*_*s*_is half of the cylindrical leg’s circumference (yellow dashed line in Fig 1F). The resistance by surface tension is higher when leg diameter, *D*, is larger:

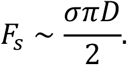

#### Total resistance on a leg

The resistance force on the leg, *R*, is the sum of *F*_*h*_, *F*_*w*_, and *F*_*s*_:

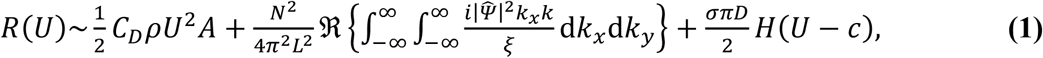

where *H*(*x*) is the Heaviside function to represent the surface tension force as a stepwise function based on velocity criteria, *c*.

#### Modeling normal forces responsible for water strider’s floating on the surface and resistance forces acting on water strider’s legs during symmetric and asymmetric sliding

Figures 1B and C show the schematics of a water strider sliding on the water surface. The normal force on a leg supporting the anterior side is *N*_*a*_ and the posterior side is *N*_*p*_ (Fig 1B). The force balance in the vertical direction is represented as

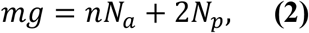

where *g* is gravitational acceleration, *m* is the mass of the water strider, and *n* is the number of legs involved in the anterior part and it depends on the sliding posture. At asymmetric sliding one midleg creating supporting force, *N*_*a*_, supports the anterior side so *n* = 1, while during symmetry sliding two fore legs, each creating supporting force, *N*_*a*_, are supporting the anterior side so *n* = 2. Hence, the total normal anterior force can be represented as: *N*_*aT*_ = *nN*_*a*_. The body is represented in the model as a uniform rod with a length corresponding to the body length of the water strider and oriented parallel to the direction of movement (this is a simplification as water striders do not keep their body axis ideally parallel to the movement direction during asymmetric sliding). The torque balance about the center of the mass of the water strider is represented as:

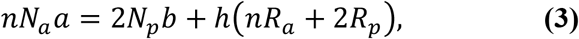

where *h* is the height of the center of the mass above the undisturbed water surface, *a* and *b* are the horizontal distances along the axis parallel to the moving direction from the center of the mass to the center of the wetted anterior supporting leg(s) and of the wetted posterior supporting leg(s), respectively. The values of *a* and *b* are calculated based on the leg segments’ lengths, the body length, the leg attachment positions on the body, and the empirically measured angles at joints between leg segments (Fig 7); *R*_*a*_ and *R*_*p*_ are the resistance forces on each of the anterior supporting legs and on each of the two posterior supporting legs, respectively, and they contribute to the total resistance (*R*_*T*_ = *nR*_*a*_ + 2*R*_*p*_).

**Fig 7.**
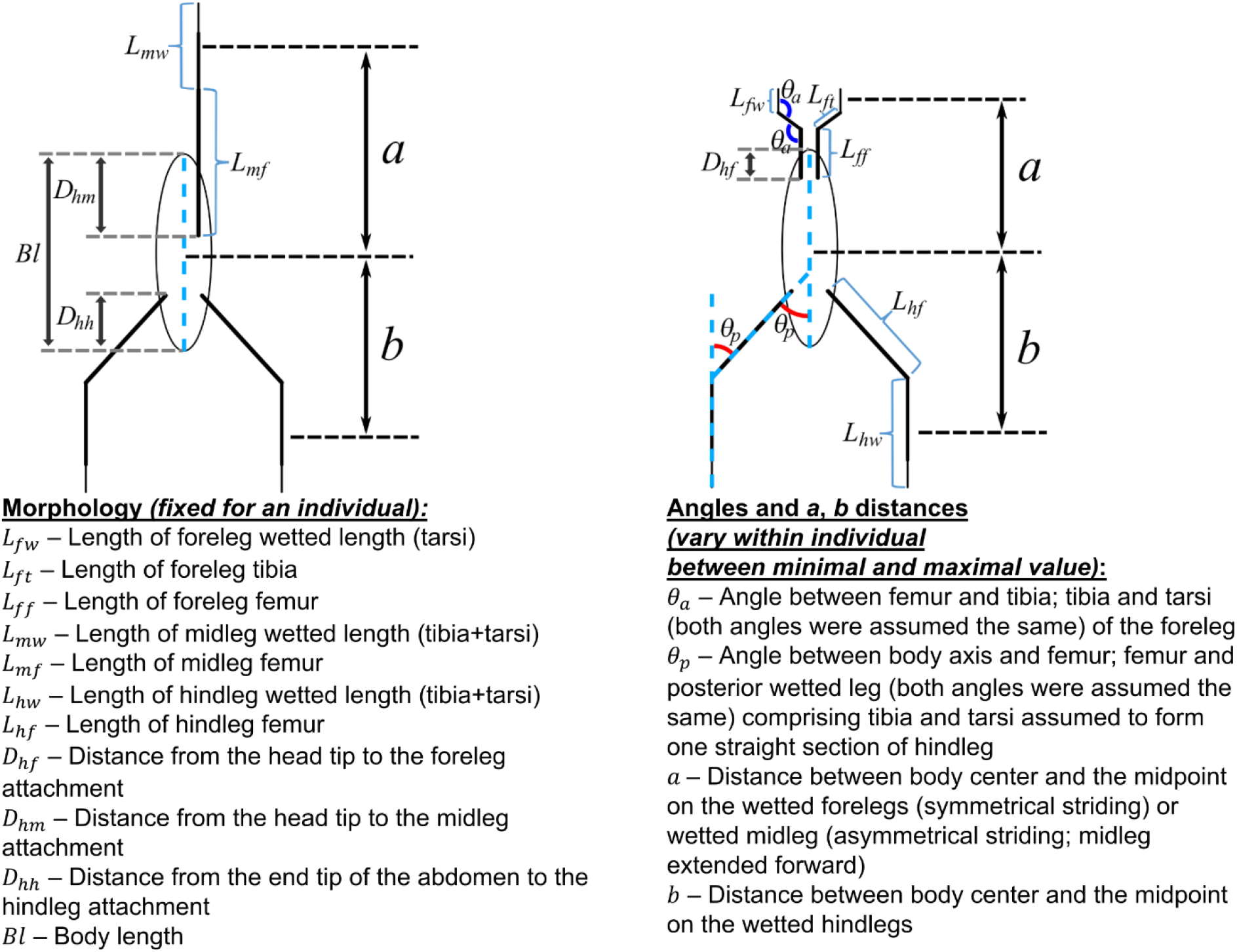
Graphical schematics of the morphological variables used in calculations in the model. See Methods for detailed explanations of how the variables were used to calculate *a* and *b*.

If predicted *N*_*a*_ is larger than the maximal surface tension force produced by anterior legs (either two forelegs in symmetrical sliding or one midleg in asymmetrical sliding), the water strider cannot float on the surface. We use the theoretical model to determine the conditions for floating on the surface (i.e., sliding without surface breaking) during symmetric and asymmetric sliding, and to calculate the resistance values for water strider’s legs that interact with water (red marked wetted legs in Fig 1A, B, C) for various sliding velocities, body mass, and leg geometries. The theoretical model reasonably well simulates the empirically observed trajectories (Fig S3).

To check if the model imitates the behavior of water striders in a reasonable manner we numerically calculated the displacement, *x*, of a sliding water strider (from the equation of motion, 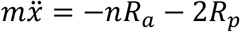, which expresses the effect of resistance on the displacement) based on empirically derived variables from two video clips, one for the *G. gigas* and one for the *A. paludum*, (Fig S3). The comparison confirmed that the theoretical model imitates reasonably well the real striding by the larger (Fig S3A) as well as the smaller (Fig S3B) water striders. We also confirmed that the shapes of the theoretically calculated curves of resistance fit reasonably well, considering the unavoidable scattering of empirical data, to the shapes of splines obtained from the generalized additive model analysis of empirical evaluations of resistance based on observed decelerations during sliding by *A. paludum* extracted from video clips (Fig S13).

### Re-distribution of leg geometry to create virtual morphologies of theoretical water strider species

To predict the model results for non-existing combinations of leg geometry, body mass and striding behavior, we chose leg geometry of *G. gigas, P. tigrina*, and *A. paludum* as representatives of “long-foreleg”, “short-foreleg”, and “intermediate-foreleg geometry”, respectively. As the leg geometries of the three small-sized species showed small differences (Fig 2), *A. paludum* was chosen as the sole representative of the “intermediate-foreleg geometry”. We used five size classes corresponding to empirically observed ranges and empirical distributions of body mass in our study species: *G. latiabdominis* (12-32 mg), *A. paludum* (35-72 mg), *P. tigrina* (83-144 mg), *G. gigas* females (217-318 mg) and *G. gigas* males (316-511 mg). We created morphological data for 15 separate situations (5 body mass classes * 3 “leg geometries”), including 5 empirically collected data (for the five size/sex classes in our study) and 10 “virtual” situations that have not been recorded in our study species. Each situation was represented by a population of individuals (actually measured or virtually created) with their morphological traits: body mass, body length, distance from the head tip to the location of foreleg attachment to the body, distance from the head tip to the location of midleg attachment to the body, distance from the head tip to the location hindleg attachment (those distances expressed as proportion of body length), and leg measurements: femur, tibia, tarsus. Wetted leg length was assumed as tarsus (forelegs) or tibia plus tarsus (midleg and hindleg). For each individual in each empirical data set we additionally expressed the femur, tibia and tarsus lengths as proportions in the total length of the legs of that individual.

We created 10 virtual data sets on the basis of the 5 empirically measured populations according to the following procedure. To create a virtual population of water striders with the distribution of body mass, and the total leg lengths observed in the sample of species *A* with the range and distribution of the “leg geometries” measured in the sample of species *B*, we redistributed the empirically measured total leg length of each of *nA* individuals of species *A* (keeping wetted leg diameter of species *A*) into the lengths of femur, tibia and tarsus of forelegs, midlegs and hindlegs according to the length proportions observed in each of *nB* individuals of species *B* (we also shifted leg attachment points relative to the body length as observed in each individual of species *B*). Hence, the virtual population comprised a total of *n*_*A*_*n*_*B*_ individuals. This process was designed to result in a conservatively wide range of estimated virtual morphologies in order to take into account all possible virtual combinations and to focus on the major differences among virtual species (differences observed even though the virtual species include extreme morphological combinations). However, for the real water striders, we only used the morphological combinations observed in nature, i.e., we did not redistribute leg lengths of one individual of species *A* into proportions observed in the remaining individuals of species *A*. In this way, we intended to consider morphological variability that actually represents the reality observed in the real water striders.

### Numerical calculations

Based on the mathematical model we built a computational model in MATLAB. At the core of the model was numerical integration of the equation of force and torque balance using morphological data of the studied species and “virtual” re-distribution. Figures showing model output were also prepared using MATLAB.

To create Fig 3A-E that maps each of the 30 different sets of water striders (either real or virtual one) representing different combinations of body mass, leg geometry and locomotion onto the phase diagram determining conditions for floating, for each individual (real or virtual) characterized by specific fixed body mass and leg morphology (Fig 7), including the value of 2(*L*_*a*_ + *D*) that comprises the horizontal axis of Fig 3A-E, we determined the minimal and the maximal value of the total anterior normal force required to support the frontal part of body during sliding on the surface (*N*_*aT*_, on the vertical axis in Fig 3A-E) based on the combination of minimal and maximal values of the following four variables that affect the normal forces (*N*_*a*_, *N*_*p*_) and resistance forces (*R*_*a*_, *R*_*p*_) in the model (in the system of equations presented in *Model description* section):

*a* - horizontal distance from the body center to the center of anterior wetted leg(s) (Fig 1B, Fig 7)

*b* - horizontal distance from the body center to the center of posterior wetted legs (Fig 1B, Fig 7)

*h*- body height above the water surface (Fig 1C)

*U* - body velocity.

In a similar manner, we determined for each individual the minimal and maximal value of total resistance *R*_*T*_ during sliding (if sliding is feasible) as a function of body velocity, *U*. The minimal and maximal values of total resistance depended on the combinations of the maximal and minimal values (for each individual) of the three remaining variables; *a, b, h*. (and the set of fixed variables for each individual such as body mass and leg morphology).

The maximal and minimal values of the total normal force (*N*_*aT*_) for each individual were marked in the phase diagram leading to an outline in Fig 3A-E for each of the 30 different sets of water striders representing different combinations of body size class, leg geometry and locomotion. The outlines of predicted total resistance (*R*_*T*_) in Fig 3F-J as a function of *U* were obtained in a similar way. We also calculated *R*_*T*_ values by using average leg morphology for each set of water striders (of given body size class, leg geometry, and locomotion mode) combined with the mid-range values of *a, b*, and *h*, (solid or dashed lines in Fig 3F-J).

For each individual (real or virtual), the ranges of *a* and *b* were calculated by using the body size and leg morphology (fixed for an individual), and the minimal and maximal values of the empirically observed values of two angles (Fig 7):

*θ*_*a*_ – angle between femur and tibia; tibia and tarsus (both angles were assumed the same) of the foreleg (in symmetrical sliding it is anterior leg - hence subscript *a*). The angle was measured in the plane common for all three sections of a leg (femur, tibia, tarsus). It was determined from photographs and videos to range from ∼90 to ∼135 degrees for all 5 size classes of water striders. These two extreme values were used to calculate the minimal and maximal values of *a* for each individual.

*θ*_*p*_ – angle between body axis and femur; femur and posterior wetted leg (both angles were assumed the same) comprising tibia and tarsus assumed to form one straight section of hindlegs. The angle was measured in the plane common for all three sections of a leg (femur, tibia, tarsus). It was determined from photographs and videos to range from ∼30 to ∼60 degrees for all 5 size classes of water striders. These two extreme values were used to calculate the minimal and maximal values of *b* for each individual.

The values of *a* and *b* were calculated by assuming that a midleg is extended forward (femur parallel to body axis/movement direction) with the wetted midleg parallel to body axis and movement direction, and that wetted hindlegs are also parallel to the body/movement axis during asymmetric sliding (left), and by assuming that tarsi and femora of forelegs, as well as hind legs’ wetted parts are always parallel to the movement axis during symmetric sliding (Fig 7).

We used the following formulas to calculate a and b.

In asymmetric sliding:

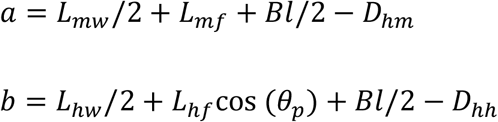

In symmetric sliding:

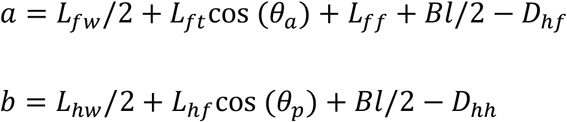

We estimated the maximal and minimal values of *h* and *U* by observations and measurements from video clips and photographs of each species. The body height ranges were determined as 3-10 mm, 3-7 mm, 1-3 mm, and 1-3 mm for *G. gigas, P. tigrina, A. paludum*, and *G. latiabdominis*, respectively. The body velocity ranges were determined as 0-1.5 m/s, 0-1.3 m/s, 0-1.2 m/s, and 0-1.0 m/s for *G. gigas, P. tigrina, A. paludum*, and *G. latiabdominis*, respectively. Wetted leg diameter, *D*, was empirically derived by using average of the five wetted leg diameters for each size category. Each wetted leg diameter of individuals was calculated by the average of the diameter of the center of wetted foreleg (tarsus), wetted midleg (tibia+tarsus), and wetted hindleg (tibia+tarsus). The wetted leg diameters, *D*, were determined as 0.266 (±0.032), 0.221 (±0.006), 0.163 (±0.017), 0.127 (±0.009), 0.077 (±0.009) mm for *G. gigas* male, *G. gigas* female, *P. tigrina, A. paludum*, and *G. latiabdominis*, respectively (average ±s.d.).

### Measurements, observations, and experiments

We determined body mass and various morphological variables explained in Fig 7 for six water strider species: *G. latiabdominis* (n=16; Seoul, Korea), *Aquarius remigis* (n=6; Huyck Preserve, USA), *G. gracilicornis* (n=16; Seoul, Korea), *A. paludum* (n=21; Seoul, Korea), *Ptilomera tigrina* (n=18; Me Linh Station for Biodiversity, Vietnam) and *Gigantometra gigas* (n=25; Pu Mat National Park, Vietnam). Photographs were used for measurements by ImageJ (https://imagej.nih.gov/ij/). The research was permitted in Pu Mat National Park by the Pu Mat National Park administration, and the study in and near the Me Linh Station for Biodiversity was permitted by the Institute of Ecology and Biological Resources, VAST, Vietnam.

We filmed *G. gigas* and *P. tigrina* in their natural habitats (standard and high-speed movies at 250, 500, and 1000 fps), and *A. paludum* and *G. latiabdominis* in acrylic containers filled with water (standard and high-speed at 1000 fps) with Sony RXIII-10 camera. A total of 50 striding events of *G. gigas* and 12 striding events by *P. tigrina* were filmed and used to determine their striding behavior, and a total of 236 striding events from 6 individuals of *A. paludum* and 13 striding behaviors from 5 individuals of *G. latiabdominis* were analyzed. The high-speed videos that were shot directly from above the water strider with scale at the level of the water surface were digitized and analyzed using Tracker program to determine the body velocity and acceleration.

For statistical comparisons of body velocity between different locomotion modes by *G. latiabdominis* (*n*=8 striding events by 4 individuals), we used Wilcoxon Signed-Rank Test (https://astatsa.com/WilcoxonTest/, https://www.aatbio.com/tools/mann-whitney-wilcoxon-signed-rank-test-calculator). For statistical comparisons of initial body velocity of sliding among three locomotion modes of *A. paludum*, (n=236 striding events from 6 individuals) we used *lmerTest* and *gamlss* packages (R version 3.6.1). The distance traveled, and the duration of sliding among three locomotion modes were also analyzed in a similar manner, but with only sliding events that were naturally ended by the water strider itself touching their midleg(s) with the water surface (e.g., excluding sliding event that ended by hitting wall, n=228 striding events from 6 individuals).

Additionally, we chose 72 striding events of *A. paludum* that have passive phase duration long enough (50-80 ms) to empirically evaluate the deceleration and subsequently the resistance. These data were analyzed using the general additive model (*gamlss* package in R). Finally, for a small subset of striding events (8 for *G. latiabdominis*, 16 for *A. paludum*, 8 for *P. tigrina*, and 5 for *G. gigas*) we digitized the striding from high-speed movies in a frame by frame manner in order to extract information for evaluation of acceleration and force generated during thrust phase of each species (seen in Fig S6).

## Acknowledgments

We would like to thank the crew of the Me Linh Station for Biodiversity. We thank the following persons who helped in various matters during our research: Dang Huy Phuong (Head of Me Linh Station for Biodiversity), Nguyen Van Mon, Nguyen Van Dat, Nguyen Van Khoi, Nguyen Van Ty, Trinh Xuan Thanh. We thank the administration of the Pu Mat National Park for permit and help in research. We thank Sohyeon Ju and Jeongwon Yoon for helping with data analysis. We thank Mihye Jun and Minkey Kim for helping with the field recording. We also thank all individuals that helped us during the field studies.

## Data and materials availability

Data sets of species measurement associated with analyses/figures are located in https://doi.org/10.5281/zenodo.6790153. The Matlab codes for the theoretical model are deposited at https://doi.org/10.5281/zenodo.6790153.

## Author contributions

Plan and design: WK, JHL, PGJ, HYK, SIL

Carry out empirical research and analysis: WK, HTP, ADT, JH, SYB, PGJ, SIL

Carry out theoretical modeling: JHL, HYK

Drafting manuscript: WK, JHL

Work on final manuscript: all co-authors

## Supporting information

Supplementary Materials.pdf – this file contains the supplementary Text, Figures S1-14, Tables S1-6, and Data with explanations.

Supplementary Movies S1 to S7

