## Supplementary materials for "Physics of sliding on water predicts morphological and behavioral allometry across a wide range of body sizes in water striders (Gerridae)"

**This PDF file includes:**

Supplementary Methods

Figures S1 to S14

Tables S1 to S6

Movies S1 to S7

**Other Supplementary Materials for this manuscript include the following:**

Movies S1 to S7

Supplementary Methods

**Additional mathematical explanations for wave drag**

*(Based mostly on Raphaël & De Gennes, 1996)-reference in the main text*

***Velocity potential***

The velocity potential of the liquid is determined by solving Laplace’s equation:

$$\nabla^{2}\varphi=0$$

with the boundary condition, $\partial\varphi/\partial z\to0$ for $z\to-\infty$. The formula of the velocity potential is

$$\varphi=\int_{-\infty}^{\infty} \int_{-\infty}^{\infty} \frac{1}{4\pi^{2}}B\left( k_{x},k_{y} \right)e^{i(k_{x}\left( x+Ut \right)+k_{y}y)}e^{kz}dk_{x}dk_{y}$$

where $k=\left( k_{x}^{2}+k_{y}^{2} \right)^{1/2}$ and $B\left( k_{x},k_{y} \right)$ is the constant. The coordinates at $t=0$ are represented in figure 1D.

The vertical displacement of liquid surface, $\zeta$, is obtained by $\partial\zeta/\partial t=\left( \partial\varphi/\partial z \right)_{z=0}$. The Fourier transform of the vertical displacement of liquid surface, $\zeta$, is $\hat{\zeta}$ and satisfies the following equation:

$$\zeta\left( x, y, t \right)=\int_{-\infty}^{\infty} \int_{-\infty}^{\infty} \frac{1}{4\pi^{2}}\hat{\zeta}\left( k_{x},k_{y} \right)e^{i(k_{x}\left( x+Ut \right)+k_{y}y)}dk_{x}dk_{y}$$

Therefore, the velocity potential is represented as

$\varphi=\int_{-\infty}^{\infty} \int_{-\infty}^{\infty} \frac{1}{4\pi^{2}}\frac{ik_{x}U}{k} \hat{\zeta}\left( k_{x},k_{y} \right)e^{i(k_{x}\left( x+Ut \right)+k_{y}y)}e^{kz}dk_{x}dk_{y}$	(S1)

The relationship of $\hat{\zeta}$ and $\hat{P}$ is determined from the Navier-Stokes equation and equation (S1):

$\left[ \rho gk+\sigma k^{3}+\rho\left( 2\nu k^{2}-ik_{x}U \right)^{2}-4\rho\nu^{2}k^{3}\sqrt{k^{2}-\frac{ik_{x}U}{\nu}} \right]\hat{\zeta}=-k\hat{P}$	(S2)

where $\hat{P}$ is the Fourier transform of the external pressure, $P\left( x,y \right)=N\left( H\left( x \right)-H\left( x-L \right) \right)\delta\left( y \right)/L$ so $\hat{P}=N(1-e^{-iLk_{x}})/{ik}_{x}L$.

***Wave drag***

The wave drag is the total pressure per unit area of the liquid surface in the x-direction:

$F_{w}=-\int_{-\infty}^{\infty} \int_{-\infty}^{\infty} P\left( x,y \right)\left( \frac{d}{dx}\zeta\left( x,y \right) \right)dxdy=-\int_{-\infty}^{\infty} \int_{-\infty}^{\infty} \frac{ik_{x}\hat{\zeta}\hat{P}}{4\pi^{2}}dk_{x}dk_{y}$	(S3)

From equation (S2) and (S3) we obtain the wave drag:

$$F_{w}=\mathfrak{R}\left\{ \int_{-\infty}^{\infty} \int_{-\infty}^{\infty} \frac{i\hat{P}^{2}k_{x}k}{4\pi^{2}\xi}dk_{x}dk_{y} \right\}=\frac{N^{2}}{4\pi^{2}L^{2}}\mathfrak{R}\left\{ \int_{-\infty}^{\infty} \int_{-\infty}^{\infty} \frac{i\left| \hat{\Psi} \right|^{2}k_{x}k}{\xi}dk_{x}dk_{y} \right\}$$

Supplementary Figures

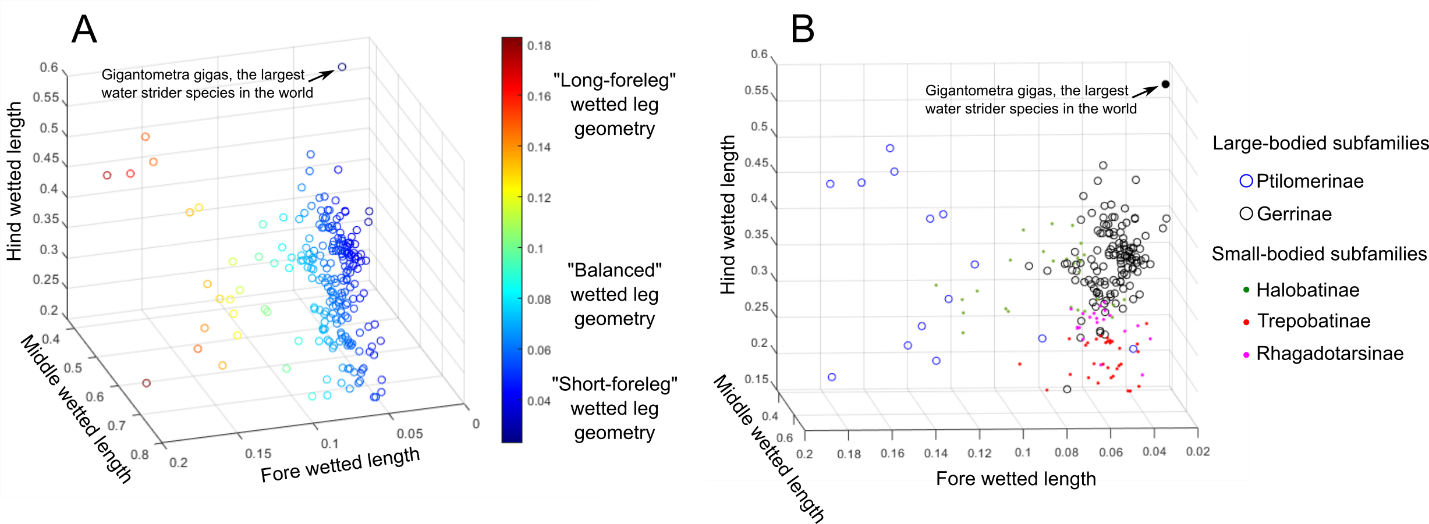
Fig S1. Distribution of wetted leg geometries among Gerridae

(A) – 3D scattergram of proportions of wetted forelegs, midlegs and hindlegs (3 axes in the plot) in the total wetted leg length with data points color coded to highlight the differences among species in the proportion of wetted forelegs; (B) – 3D scattergram similar to (A) with data points marking different subfamilies (according to Matsuda 1960) and family-typical body size of a water strider (families with the typically larger size species are marked with color-coded unfilled circles and families with the typically smaller size species are marked with color-coded dots. Figure is based on Table 16 in Matsuda 1960. The concept of the “wetted leg geometry” is crucial in our theoretical model. The term refers to the relative proportions of wetted forelegs, wetted midlegs and wetted hindlegs in the total length of the wetted legs. The figure suggests that we can classify species into at least three types of “wetted leg geometry”: the “intermediate-foreleg” or “standard” observed in the frequently studied small and mid-size genera *Gerris* and *Aquarius*, (wetted forelegs comprise from ~4 to ~8% of the total wetted legs length), the “long-foreleg geometry” known in the marine small water striders, Halobatinae, and in the medium and large water striders from the subfamily Ptilomerinae (wetted forelegs comprise about 18% of the total wetted legs length), and the “short-foreleg geometry” (wetted forelegs comprise less than ~4% of the total wetted legs length) well documented in at least one large species, *G. gigas,* (Tseng and Rowe 1999), and present in some larger species of Gerrinae.

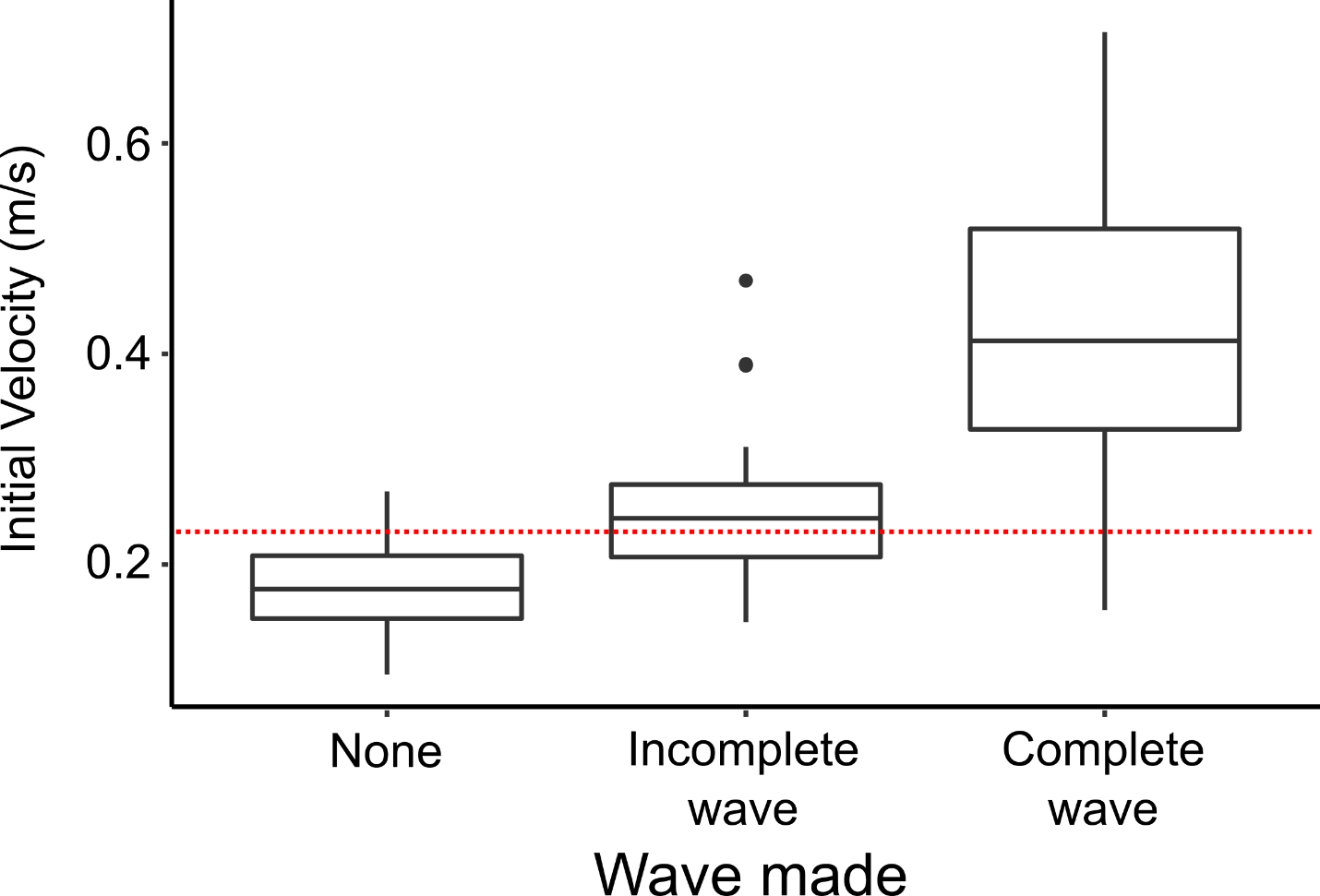

Fig S2. Empirical validation of the assumptions about velocity that leads to capillary gravity waves that increase resistance

The minimum critical theoretical velocity resulting in capillary gravity waves, $\boldsymbol{c}$, is 0.231 m/s (marked with the red doted horizontal line). Comparison of water strider *A. paludum* initial sliding velocity among sliding without visible waves, sliding with incomplete/weak waves and sliding with clearly visible waves shows that waves start being noticeable around the velocities similar to the critical velocity.

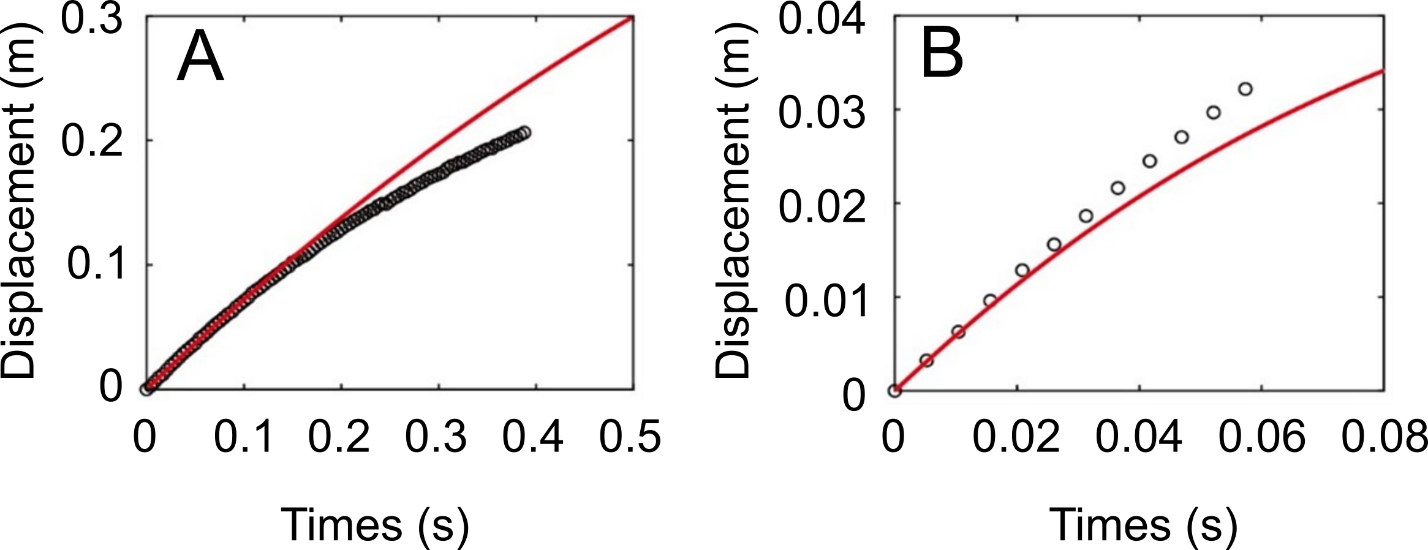
Fig S3. Validation of the theoretical model accurateness in imitating sliding of larger and smaller species of Gerridae

(A) Empirical displacement of the larger species (*G. gigas*) sliding asymmetrically. (B) Empirical displacement of the smaller species (*A. paludum*) sliding symmetrically. The red solid line corresponds to theoretical predictions and the circles correspond to empirical data extracted from two movie clips of sliding.

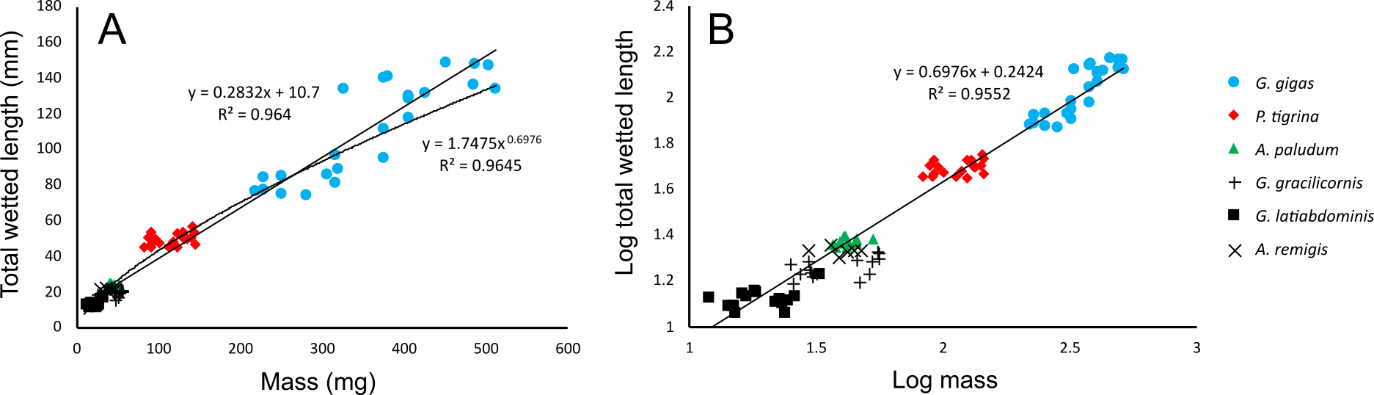

Fig S4. Relationship between the total wetted length and body mass of the six studied species

Body mass and total wetted length are marked for *Gigantometra gigas* (blue circles), *P. tigrina* (red diamonds), *Aquarius paludum* (green triangles), *Gerris gracilicornis* (plus signs), *G. latiabdominis* (black squares), and *A. remigis* (cross-marks) in normal (A) and log scale (B).

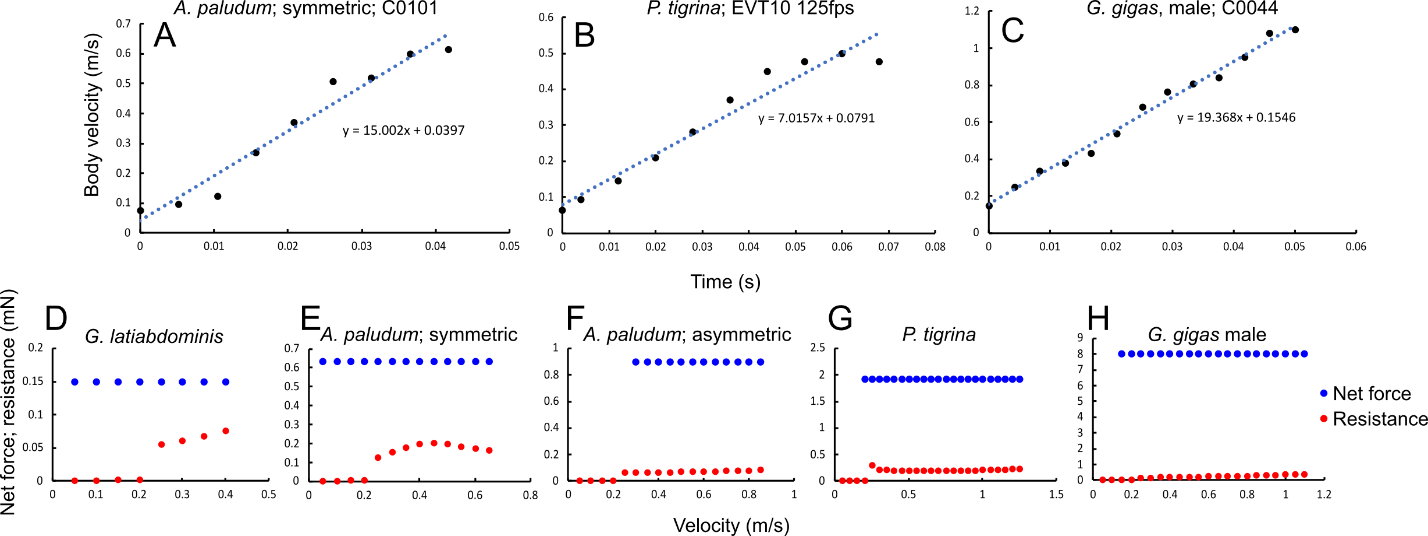

Fig S5. Three examples of digitized thrusting phase of a stride

Strides to explain the method of extracting the thrusting phase acceleration as a slope of a regression line fitted to the data on body velocity versus time for each stride, and five examples of strides for which the net thrust force (based on the acceleration and body mass: $\boldsymbol{m*a}$) and the theoretically predicted resistance force are depicted. (A) – a thrusting phase of a symmetric stride of *Aquarius paludum*; (B) – a thrusting phase of a symmetric stride of *Ptilomera tigrina*; (C) – a thrusting phase of an asymmetric stride of *Gigantometra gigas* male. These estimated accelerations for each stride of an individual of known body mass were used to calculate the thrust net force ($\boldsymbol{F=ma}$) in each stride. Thrusting phase is defined as the duration from the first frame with leg pushing backwards till the frame with maximum body velocity; it varies among species as the range of velocity (m/s) on x-axis in d-h show. (D-H) – examples of force profiles during thrust phase where the empirically evaluated thrust net force (blue dots) and the theoretically predicted resistance (red dots) force (based on empirically measured body velocity profile) are displayed in single strides of *Gerris latiabdominis* (d; symmetrical stride), *A. paludum* (symmetrical stride in e and asymmetrical stride in f), *P. tigrina* (G) and *G. gigas* (H).

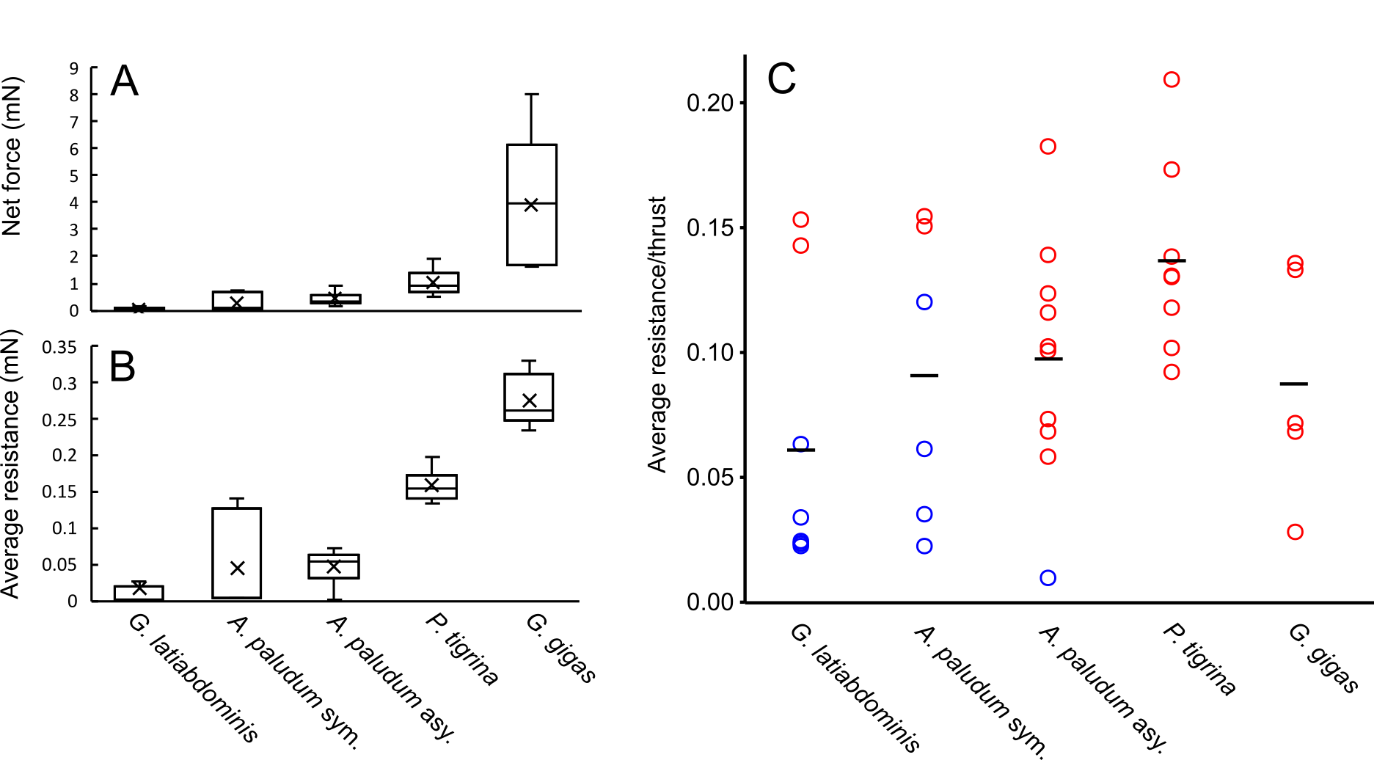

Fig S6. Empirical net force and theoretical resistance for each species

Comparison of empirically derived (see Fig S5) the net thrust force (A), which directly contributes to the momentum change of insect body during thrust phase of the striding, the theoretically predicted resistance force (B), which has to be countered by the insect using the thrust. (C) The proportion of the resistance in the total thrust produced by an insect (resistance / (net force +resistance)) in the chosen sets of strides in the four study species: *Gerris latiabdominis* striding symmetrically (n=8), *Aquarius paludum* striding symmetrically (n=6), *A. paludum* striding asymmetrically (n=10), *Ptilomera tigrina* striding symmetrically (n=8) and *Gigantometra gigas* striding asymmetrically (n=5). The average values for each species are marked as horizontal lines in (C). Red circles indicate striding at the body velocities that are higher than the theoretical body velocity threshold associated with surface waves (0.2313 m/s).

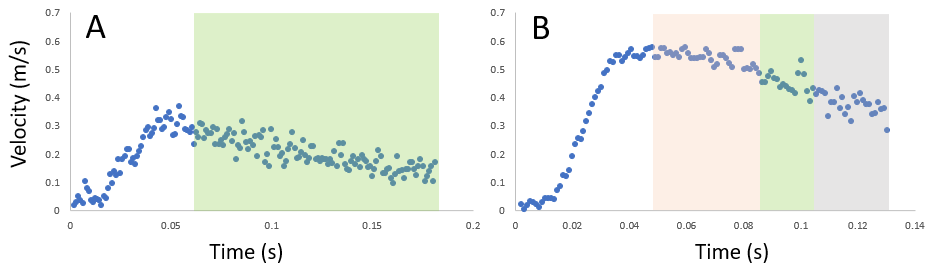
Fig S7. Symmetric sliding and leaping of *G. latiabdominis*

An example of symmetric sliding (A) and leaping (B) of *G. latiabdominis* Individual 1 (19.1 mg). Orange, green, and gray shading represent leaping, symmetric sliding, and resting phase, respectively.

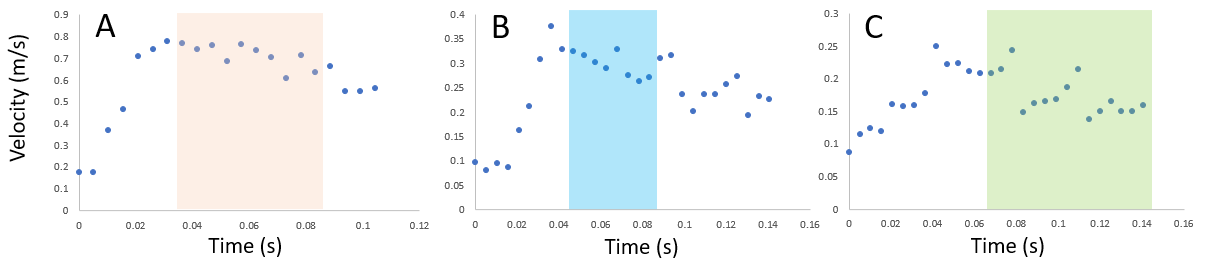

Fig S8. Symmetric/asymmetric sliding and leaping of *A. paludum*

An example of leaping (A), asymmetric sliding (B), and symmetric sliding (C) of *A. paludum* Individual 4093 (18 mg). Orange, blue, and green shading represent leaping, asymmetric sliding, and symmetric sliding, respectively.

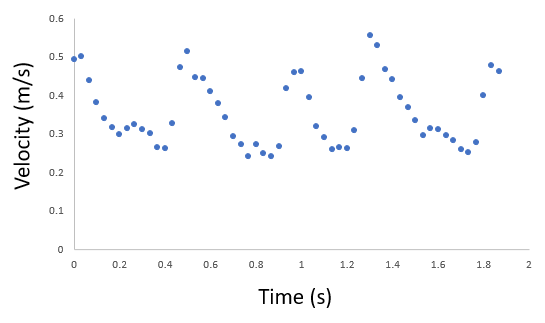

**Fig S9. Series of striding of *P. tigrina* in the field**

An example of the changes of the body (body center) velocity relative to the water surface flow in a natural situation of multiple striding by *P. tigrina* (the flow speed at the water surface is considered in the calculations of the body speed relative to the water surface).

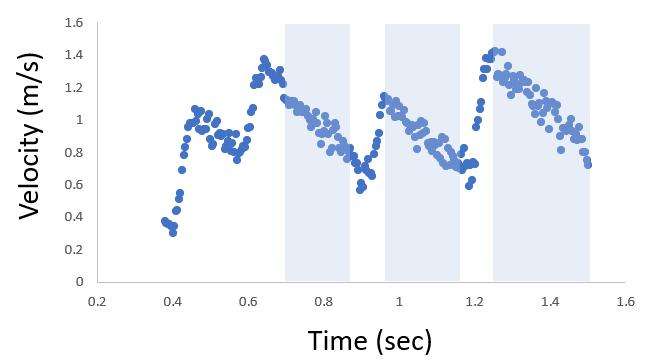
Fig S10. Series of striding of *G. gigas* in the field

An example of the changes of the body (body center) velocity relative to the water surface flow in a natural situation of multiple striding by *G. gigas* (the flow speed at the water surface is considered in the calculations of the body speed relative to the water surface).

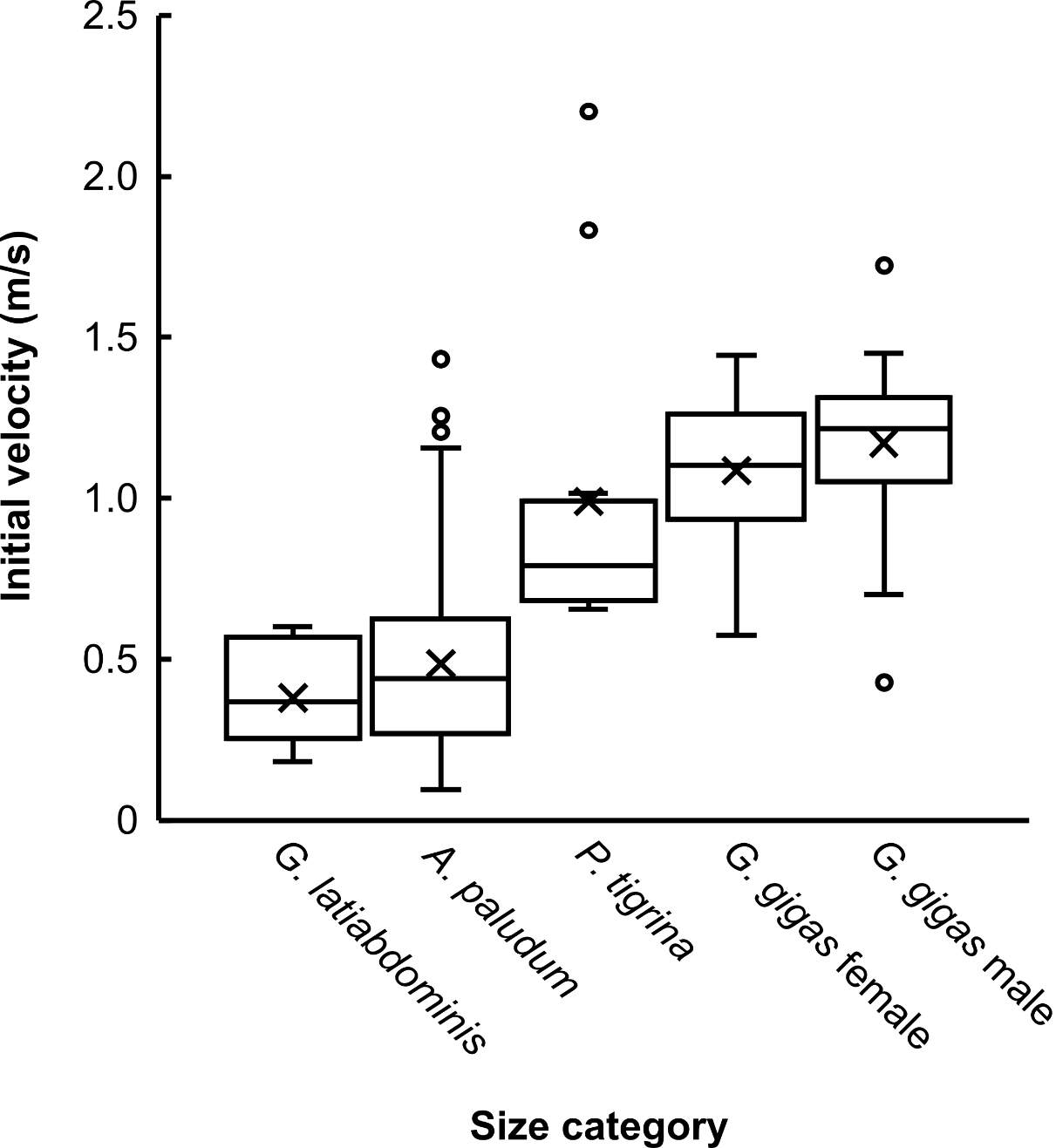

Fig S11. Box-and-whiskers plot of initial striding velocity five size classes corresponding to five species/sex classes of study subjects

*G. latiabdominis* (n=13), *A. paludum* (n=236), *P. tigrina* (n=12), *G. gigas* females (n=23), and *G. gigas* males (n=27). For *G. latiabdominis*, velocities of symmetrical sliding (n=9) and leaping (n=4) are pooled. For *A. paludum*, velocities of asymmetrical sliding (n=136), symmetrical sliding (n=68), and leaping (n=32) are pooled. Initial velocity was chosen as a peak velocity before sliding in *G. latiabdominis*, *P. tigrina*, and *G. gigas* (See methods section for *A. paludum*).

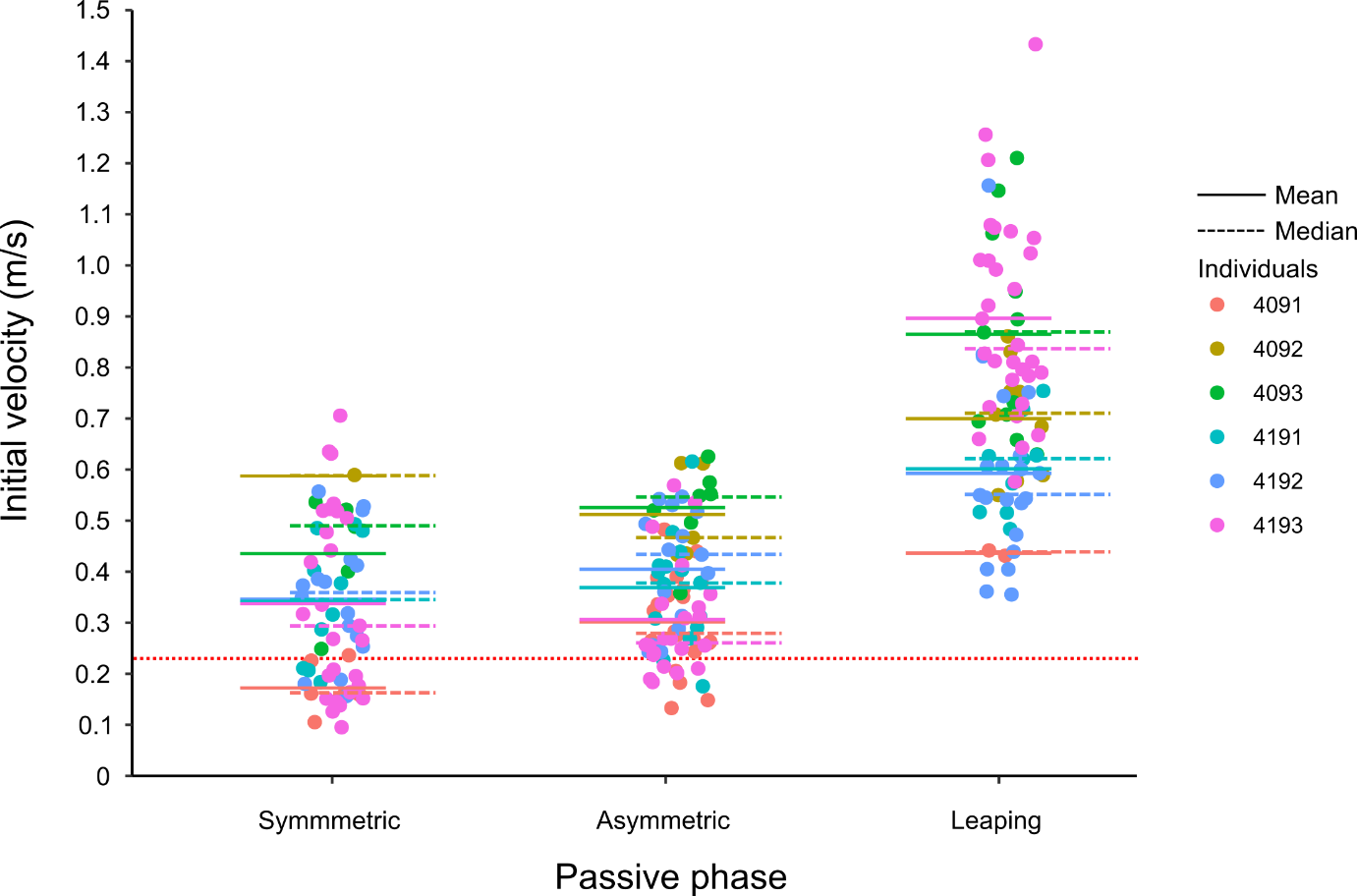

Fig S12. Initial velocity of passive phase for each individual.

The scatter plot of initial velocity in a passive phase from empirical data of *A. paludum*. Mean and average values are marked as solid and dashed lines, respectively. The red dotted line represents wave making velocity (0.23 m/s). The statistical analysis results are in Table S3.

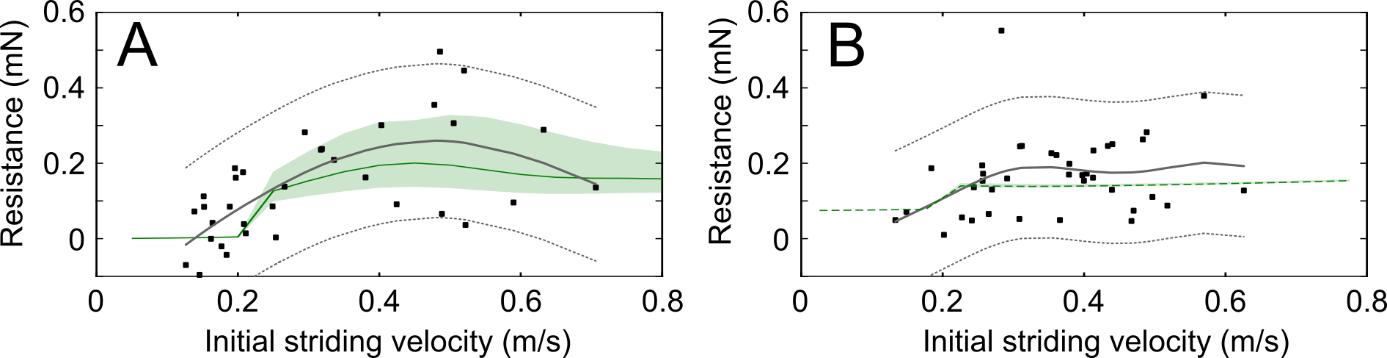
Fig S13. Resistance for *A. paludum*’s sliding from mathematical model and from empirical calculations.

Results of the model calculation and empirical analysis for symmetric (A) and asymmetric (B) sliding of *A. paludum*. Theoretical calculations from mathematical model are marked as green line and shadings. Black squares represent empirical data points, the generalized additive models from those data are marked as gray solid line, and 97.5% centiles are also marked with dashed lines. Resistance was calculated from the estimated deceleration (Table S6 on the next two pages) and body mass for each stride. The theoretical predictions in this figure were also used in Fig 5D of the main text.

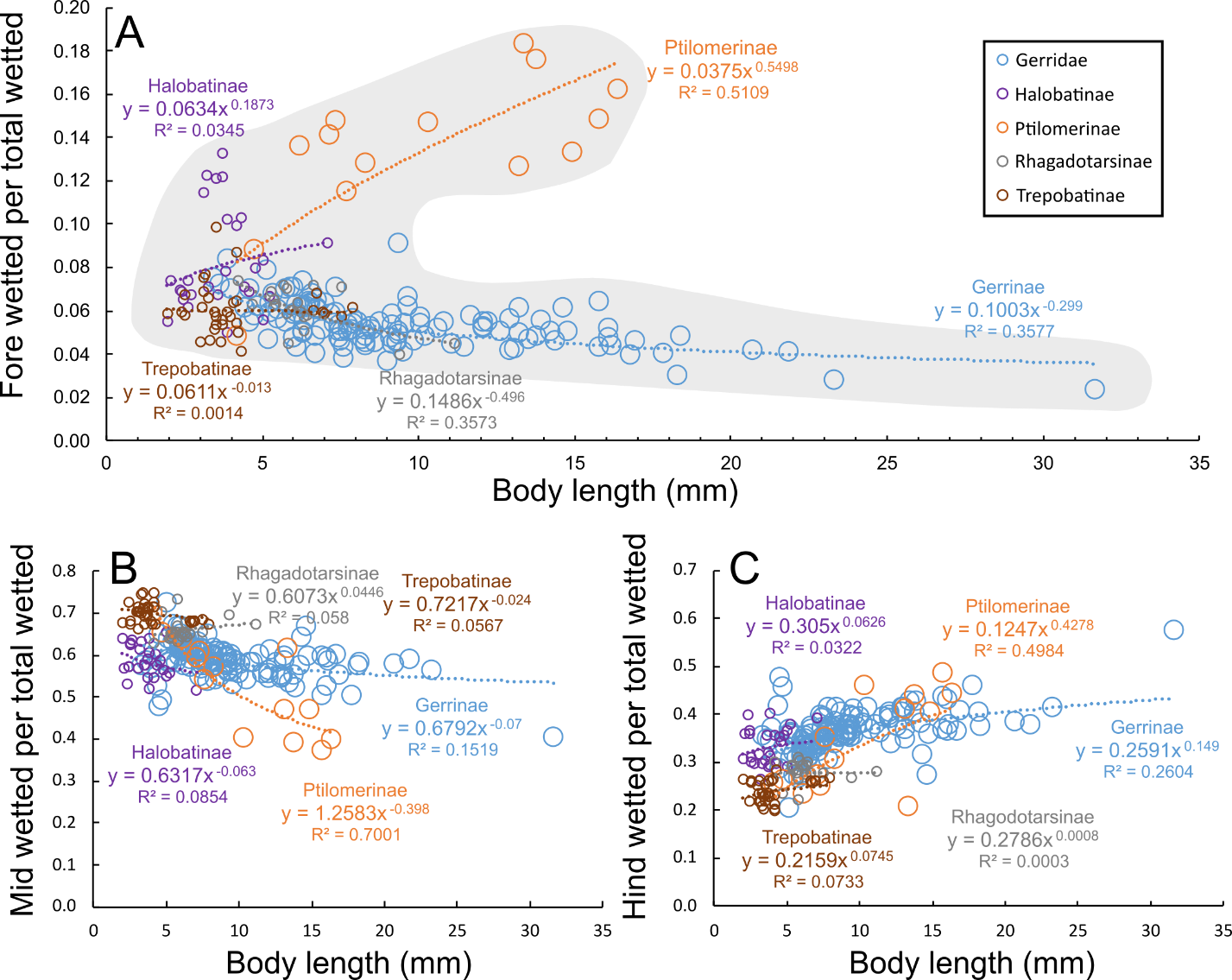

**Fig S14.** **Relationship between the body length and the “wetted leg geometry”** Relationship between the body length and the proportion of wetted forelegs (A), wetted midlegs (B) and wetted hindlegs (C) in total wetted legs length. The gray shaded area in (A) helps visualizing that with increasing body size the water striders adopt one of two “wetted leg geometries”, either “long-foreleg” or “short-foreleg geometry”. The species are represented by subfamilies: *Gerrinae* (blue circles), *Ptilomerinae* (orange circles), *Halobatinae* (purple circles), *Rhagadotarsinae* (gray circles), and *Trepobatinae* (brown circles). The data re from Table 16 in Matsuda 1969, and the subfamilies follow Matsuda 1960, which may be not entirely consistent with the more modern assignments of genera into subfamilies. Additionally, this is phylogenetically un-corrected relationship, and therefore it does not directly represent evolutionary processes shaping the evolutionary changes of leg morphology as a function of evolutionary changes of body size. The equations fitted to the data points for each family separately have the power form following the convention for allometric equations. However, we used body length because of the absence of data for body mass, absence of body width, height or diameter data, and absence of body length – body mass formulas for water striders over such a large body size range. We expect that from among the possible linear measurements of body (width, height, length) the length is relatively more correlated with the body mass (albeit not necessarily in a linear fashion) than are body width or height as they are relatively small and differ among species relatively less than the body length. We decided not to use *body length^3^* (a possible alternative used occasionally in allometry) because of the elongated shape of the water striders. For the overview of leg geometries in Gerridae of various subfamilies and various body sizes we used data from Table 16 in Matsuda (1960; see also Fig 6 based on the same data). We converted the Matsuda’s units to metric measurement units (mm) using the information provided in Matsuda (1960) after one correction that was necessary to circumvent Matsuda’s apparent mistake. On page 32 Matsuda states the following rules: “*In table 16, 82 units are equal to 10 mm. For those values with asterisks, 173.7 units are equal to 10 mm*.”. However, we concluded that the correct rule is “10 mm is equal to the 173.7 units of measurements presented in Table 16” for all species listed there (regardless of whether they are marked with asterisk or not) because only this rule gives results that agree with body lengths of several species known to us, and with the body length of *G. gigas* listed by Matsuda in a different part of his book. Matsuda mentions on page 12 that that body length of *G. gigas* is 3.19 cm (“*Gigantometra gigas* (China), male. Length of body: 31.9 mm.”; Fig 3), which is consistent with the calculation using the “173.7 units” rule but not the” 82 units rule” (predicted body length of G. gigas: 67.08 mm), which according to Matsuda’s erroneous advice is supposed to be applied to *G. gigas* as it is listed without an asterisk in Table 16. When we tried to convert body length by using the “173.7 units rule” to species without asterisks then the body lengths matched reasonably well with our observed data (e.g., ~31 mm for *G. gigas*, ~12 mm for *A. paludum*, ~13-16 mm for Ptilomerinae). Therefore, we used the “173.7 units rule” for data presented in Figures 6 and this figure.

Supplementary Tables

Table S1. Explanations of the symbols in the model

The lists of symbols used in the mathematical model. The symbols below black solid line used only in supplementary materials. Values used in the model are shown for variables that do not change values among different simulations. The symbols are listed in the order in which they appear in the text.

| Explanations of the symbols in the model | |
| --- | --- |
| $R$ | Resistance force on a leg |
| $F_{h}$ | Hydrodynamic drag on a leg |
| $F_{w}$ | Wave drag on a leg |
| $F_{s}$ | Surface tension force on a leg |
| $\rho$ | Density of water, 997 kg/m^3^ |
| $U$ | Velocity of sliding object on the water surface |
| $D$ | Diameter of a wetted leg |
| $L$ | Length of a wetted leg |
| $A=\pi DL/2$ | Wetted area assumed as a half of curved surface of a cylinder with a length, $L$, and diameter, $D$. |
| $\nu$ | Kinematic viscosity of water, 1.003 mm^2^/s |
| $C_{D}=1.328\sqrt{\nu/(UL)}$ | Drag coefficient on the flat plate |
| $\sigma$ | Surface tension coefficient of water, 0.0728 N/m |
| $g$ | Gravitational acceleration, 9.8 m/s^2^ |
| $c=\left( 4g\sigma/\rho\right)^{1/4}$ | The theoretical minimum critical velocity of a floating object on the water surface, to produces capillary-gravity |
| $N$ | Normal force on a leg from the water |
| ${k=\left( k_{x}^{2}+k_{y}^{2} \right)}^{1/2}$ | Wave number |
| $\Psi$ | Shape of wetted area; the shape of the wetted leg assumed as a line with length $L$ with the longitudinal movement |
| $\hat{\Psi}=(1-e^{-iLk_{x}})/{ik}_{x}$ | Fourier transform of $\Psi$ |
| $H(x)$ | The Heaviside function |
| $m$ | Mass of a water strider |
| $n$ | The number of the anterior supporting legs |
| $N_{a}$ | Normal force (perpendicular to the water surface) on an anterior leg: a wetted foreleg for symmetrical sliding; a wetted midleg for asymmetrical sliding |
| $N_{p}$ | Normal force on a posterior leg, i.e., on the wetted hindleg |
| $N_{aT}$ | Normal anterior force on all anterior legs (total normal anterior force) |
| $R_{a}$ | Resistance force on an anterior leg: a wetted foreleg for symmetrical sliding; a wetted midleg for asymmetrical sliding. |
| $R_{p}$ | Resistance force on a posterior leg, i.e., on the wetted hindleg |
| $h$ | Height; the vertical distance between the center of the mass and the water surface |
| $a$ | Horizontal distance in the parallel axis to the moving direction from the center of the mass to the center of the wetted anterior supporting leg |
| $b$ | Horizontal distance in the parallel axis to the moving direction from the center of the mass to the center of the wetted posterior supporting leg |
| $R_{T}$ | Resistance on all legs (total resistance, $nR_{a}+2R_{p}$) |
| $x$ | Displacement of a water strider by sliding |
| $\ddot{x}$ | Acceleration/deceleration of a water strider |
| $L_{a}$ | A wetted length of all anterior leg(s) (total anterior wetted length) |
| $L_{fw}$ | Length of foreleg wetted length (tarsus) |
| $L_{ft}$ | Length of foreleg tibia |
| $L_{ff}$ | Length of foreleg femur |
| $L_{mw}$ | Length of midleg wetted length (tibia+tarsus) |
| $L_{mf}$ | Length of midleg femur |
| $L_{hw}$ | Length of hindleg wetted length (tibia+tarsus) |
| $L_{hf}$ | Length of hindleg femur |
| $D_{hf}$ | Distance from the head tip to the foreleg attachment |
| $D_{hm}$ | Distance from the head tip to the midleg attachment |
| $D_{hh}$ | Distance from the end tip of the abdomen to the hindleg attachment |
| $Bl$ | Body length |
| $\theta_{a}$ | Angle between femur and tibia; tibia and tarsus (both angles were assumed the same) of the foreleg |
| $\theta_{p}$ | Angle between body axis and femur; femur and posterior wetted leg (both angles were assumed the same) comprising tibia and tarsus assumed to form one straight section of hindleg |
| $\varphi$ | Velocity potential |
| $B\left( k_{x},k_{y} \right)$ | Amplitude of the waves of the disturbance |
| $t$ | Time |
| $\zeta$ | Vertical displacement of liquid surface |
| $P$ | External pressure |
| $\delta$ | The Dirac $\delta$ function |

Table S2. Initial velocity for locomotion type of *G. latiabdominis*

Results The initial velocities were significantly different between sliding and leaping of *G. latiabdominis* (Wilcoxon Signed-Rank Test, p-value = 0.029).

| Individual | Video | Locomotion | Initial velocity (m/s) |
| --- | --- | --- | --- |
| 1 | C0030 | Sliding | 0.282334 |
| 2 | C0040 | Sliding | 0.373578 |
| 3 | C0053 | Sliding | 0.209729 |
| 5 | C0069 | Sliding | 0.191976 |
| 1 | C0035 | Leaping | 0.574271 |
| 2 | C0042 | Leaping | 0.596126 |
| 3 | C0057 | Leaping | 0.506352 |
| 5 | C0073 | Leaping | 0.390899 |

Table S3. Linear mixed model of initial velocity and sliding type of *A. paludum*

Results of the linear mixed model: Velocity ~ (Sliding type) + (1 | Individual) + (1 | Video:Individual), No. of observations: 236, No. of Individuals: 6, No. of random factor (Video:Individual) groups: 100. The reference passive phase type is Asymmetry. Results are shown in Fig 5A.

|  | Estimate | df | t value | Pr(>\|t\|) |
| --- | --- | --- | --- | --- |
| (Intercept) | 0.43565 | 5.40296 | 10.495 | 8.51e-05 |
| Leaping | 0.24132 | 188.12922 | 10.857 | < 2e-16 |
| Symmetry | -0.07078 | 159.28365 | -3.525 | 5.54e-04 |

**Table S4. Generalized mixed model of sliding distance and sliding type of *A. paludum***

Results of the generalized mixed model: Sliding distance ~ (Sliding type) + (1 | Individual) + (1 | Video:Individual), Family: ("Inverse Gamma"), No. of observations: 228, No. of Individuals: 6, No. of random factor (Video:Individual) groups: 100, Degrees of Freedom for the fit: 48.63265, Residual Deg. of Freedom: 179.3673. The reference passive phase type is Leaping. $E(distance)=sqrt\left( \exp\left( \sigma^{2} \right) \right)*\exp(\mu)$. Results are shown in Fig 5B.

| $\mu$ coefficient | Estimate | t value | Pr(>\|t\|) |
| --- | --- | --- | --- |
| (Intercept) | 2.72043 | 57.657 | < 2e-16 |
| Asymmetry | 0.23134 | 3.665 | 3.25e-04 |
| Symmetry | -0.38289 | -5.703 | 4.77e-08 |
| $\sigma$ coefficient | Estimate | t value | Pr(>\|t\|) |
| (Intercept) | -0.91275 | -20 | < 2e-16 |

Table S5. Generalized mixed model of sliding duration and sliding type of *A. paludum* Results of the generalized mixed model: Sliding duration ~ (Sliding type) + (1 | Individual) + (1 | Video:Individual), Family: ("Box-Cox-Cole-Green"), No. of observations: 228, No. of Individuals: 6, No. of random factor (Video:Individual) groups: 100, Degrees of Freedom for the fit: 52.71993, Residual Deg. of Freedom: 175.2801. The reference passive phase type is Symmetry. Results are shown in Fig 5C.

|  | Estimate | t value | Pr(>\|t\|) |
| --- | --- | --- | --- |
| (Intercept) | 43.907 | 26.799 | < 2e-16 |
| Asymmetry | 37.615 | 11.913 | < 2e-16 |
| Leaping | -15.615 | -9.218 | < 2e-16 |

Table S6. Deceleration data of the sliding of *A. paludum*

Sliding events that have enough passive phase duration (50-80 ms) to calculate deceleration were chosen.

| Individual | Mass  (mg) | Video | Striding  type | Initial  velocity (m/s) | Sliding/leaping  distance (mm) | Duration  (ms) | Last  velocity (m/s) |
| --- | --- | --- | --- | --- | --- | --- | --- |
| 4091 | 53 | C0082 | Asy. | 0.283 | 6.432 | 45.879 | 0.077 |
| 4091 | 53 | C0076 | Asy. | 0.133 | 7.792 | 60.477 | 0.133 |
| 4091 | 53 | C0082 | Asy. | 0.148 | 4.812 | 32.324 | 0.139 |
| 4193 | 47 | C0176 | Asy. | 0.257 | 14.146 | 66.733 | 0.143 |
| 4193 | 47 | C0162 | Asy. | 0.184 | 10.036 | 64.648 | 0.149 |
| 4193 | 47 | C0158 | Asy. | 0.309 | 16.721 | 66.733 | 0.167 |
| 4193 | 47 | C0173 | Asy. | 0.201 | 12.991 | 72.99 | 0.176 |
| 4192 | 32 | C0144 | Asy. | 0.256 | 11.019 | 45.879 | 0.179 |
| 4191 | 40 | C0114 | Asy. | 0.227 | 12.461 | 56.306 | 0.185 |
| 4191 | 40 | C0113 | Asy. | 0.291 | 13.867 | 54.221 | 0.191 |
| 4193 | 47 | C0158 | Asy. | 0.256 | 16.158 | 71.947 | 0.192 |
| 4192 | 32 | C0144 | Asy. | 0.244 | 10.768 | 44.837 | 0.196 |
| 4191 | 40 | C0114 | Asy. | 0.312 | 15.115 | 55.264 | 0.197 |
| 4193 | 47 | C0153 | Asy. | 0.413 | 23.942 | 68.819 | 0.211 |
| 4091 | 53 | C0080 | Asy. | 0.265 | 16.433 | 58.392 | 0.213 |
| 4192 | 32 | C0144 | Asy. | 0.361 | 18.669 | 65.691 | 0.225 |
| 4193 | 47 | C0149 | Asy. | 0.241 | 17.414 | 68.819 | 0.227 |
| 4091 | 53 | C0082 | Asy. | 0.379 | 22.887 | 63.605 | 0.244 |
| 4191 | 40 | C0107 | Asy. | 0.269 | 13.093 | 43.794 | 0.245 |
| 4191 | 40 | C0107 | Asy. | 0.403 | 25.87 | 71.947 | 0.256 |
| 4191 | 40 | C0108 | Asy. | 0.308 | 13.699 | 39.623 | 0.261 |
| 4091 | 53 | C0081 | Asy. | 0.353 | 18.556 | 56.306 | 0.264 |
| 4191 | 40 | C0105 | Asy. | 0.412 | 19.452 | 46.922 | 0.266 |
| 4091 | 53 | C0078 | Asy. | 0.44 | 27.142 | 70.904 | 0.278 |
| 4192 | 32 | C0142 | Asy. | 0.47 | 28.366 | 60.477 | 0.287 |
| 4191 | 40 | C0106 | Asy. | 0.378 | 17.043 | 43.794 | 0.288 |
| 4193 | 47 | C0155 | Asy. | 0.569 | 30.39 | 67.776 | 0.3 |
| 4192 | 32 | C0134 | Asy. | 0.397 | 15.897 | 33.367 | 0.303 |
| 4191 | 40 | C0111 | Asy. | 0.399 | 25.955 | 62.563 | 0.307 |
| 4091 | 53 | C0076 | Asy. | 0.365 | 23.597 | 57.349 | 0.313 |
| 4093 | 18 | C0099 | Asy. | 0.496 | 30.037 | 64.648 | 0.316 |
| 4193 | 47 | C0153 | Asy. | 0.488 | 28.948 | 64.648 | 0.316 |
| 4192 | 32 | C0123 | Asy. | 0.517 | 26.537 | 52.135 | 0.331 |
| 4191 | 40 | C0121 | Asy. | 0.439 | 23.082 | 47.965 | 0.348 |
| 4091 | 53 | C0078 | Asy. | 0.483 | 23.584 | 46.922 | 0.361 |
| 4092 | 47 | C0092 | Asy. | 0.467 | 26.894 | 54.221 | 0.375 |
| 4092 | 47 | C0093 | Asy. | 0.433 | 25.654 | 59.434 | 0.387 |
| 4093 | 18 | C0099 | Asy. | 0.625 | 34.342 | 59.434 | 0.418 |
| 4193 | 47 | C0172 | Sym. | 0.196 | 8.823 | 69.862 | 0.061 |
| 4193 | 47 | C0148 | Sym. | 0.138 | 5.533 | 55.264 | 0.082 |
| 4193 | 47 | C0158 | Sym. | 0.151 | 7.782 | 51.093 | 0.123 |
| 4193 | 47 | C0162 | Sym. | 0.177 | 9.598 | 47.965 | 0.129 |
| 4191 | 40 | C0113 | Sym. | 0.207 | 9.132 | 49.007 | 0.134 |
| 4192 | 32 | C0134 | Sym. | 0.318 | 13.373 | 57.349 | 0.136 |
| 4193 | 47 | C0162 | Sym. | 0.126 | 8.748 | 52.135 | 0.136 |
| 4193 | 47 | C0151 | Sym. | 0.164 | 9.241 | 54.221 | 0.15 |
| 4193 | 47 | C0162 | Sym. | 0.197 | 8.095 | 38.58 | 0.155 |
| 4193 | 47 | C0151 | Sym. | 0.152 | 8.343 | 45.879 | 0.157 |
| 4091 | 53 | C0079 | Sym. | 0.145 | 8.076 | 39.623 | 0.157 |
| 4193 | 47 | C0149 | Sym. | 0.317 | 14.846 | 58.392 | 0.158 |
| 4192 | 32 | C0144 | Sym. | 0.188 | 10.622 | 52.135 | 0.16 |
| 4192 | 32 | C0135 | Sym. | 0.253 | 16.521 | 66.733 | 0.161 |
| 4093 | 18 | C0095 | Sym. | 0.249 | 14.698 | 69.862 | 0.163 |
| 4193 | 47 | C0156 | Sym. | 0.266 | 14.017 | 54.221 | 0.171 |
| 4193 | 47 | C0155 | Sym. | 0.208 | 13.218 | 64.648 | 0.174 |
| 4191 | 40 | C0120 | Sym. | 0.211 | 8.902 | 34.409 | 0.175 |
| 4091 | 53 | C0077 | Sym. | 0.162 | 7.8 | 36.495 | 0.192 |
| 4191 | 40 | C0116 | Sym. | 0.184 | 8.439 | 32.324 | 0.207 |
| 4193 | 47 | C0149 | Sym. | 0.294 | 16.8 | 64.648 | 0.215 |
| 4191 | 40 | C0121 | Sym. | 0.485 | 13.711 | 31.281 | 0.227 |
| 4191 | 40 | C0121 | Sym. | 0.403 | 13.998 | 31.281 | 0.246 |
| 4192 | 32 | C0126 | Sym. | 0.38 | 18.202 | 42.751 | 0.251 |
| 4193 | 47 | C0178 | Sym. | 0.336 | 16.381 | 47.965 | 0.27 |
| 4193 | 47 | C0173 | Sym. | 0.519 | 29.176 | 66.733 | 0.274 |
| 4193 | 47 | C0178 | Sym. | 0.477 | 24.928 | 56.306 | 0.294 |
| 4192 | 32 | C0139 | Sym. | 0.424 | 19.841 | 42.751 | 0.309 |
| 4093 | 18 | C0101 | Sym. | 0.488 | 25.563 | 47.965 | 0.348 |
| 4193 | 47 | C0176 | Sym. | 0.505 | 26.002 | 52.135 | 0.359 |
| 4193 | 47 | C0170 | Sym. | 0.632 | 35.399 | 59.434 | 0.422 |
| 4093 | 18 | C0100 | Sym. | 0.521 | 24.456 | 37.538 | 0.456 |
| 4092 | 47 | C0092 | Sym. | 0.589 | 30.869 | 45.879 | 0.483 |
| 4193 | 47 | C0164 | Sym. | 0.706 | 35.184 | 45.879 | 0.536 |

**Supplementary Movies**

**Movie S1. *G. latiabdominis* striding in the water container**

1-10 s. – symmetric striding, slowed down 16x; 11-17s. – leaping, slowed down 16x.

**Movie S2. *A. paludum* striding in the field and the water container**

1-6 s. – symmetric striding in the field, slowed down 8x; 7-9s. – symmetric striding in the water container, slowed down 8x; 10-17 s. – asymmetric striding in the field, slowed down 8x; 18-21 s. – asymmetric striding in the water container, slowed down 8x; 22-24 s. – leaping in the field, slowed down 8x; 25-28 s. – leaping in the water container, slowed down 8x.

**Movie S3. *P. tigrina* striding in the field**

1-7 s. – symmetric striding in the field, play speed 1x; 8-35 s. – symmetric striding in the field, slowed down 16x.

**Movie S4. *P. tigrina* grooming in the water-flowing container**

1-28 s. – asymmetric striding while grooming in the water-flowing container, slowed down 16x.

**Movie S5. *G. gigas* striding in the field**

1-9 s. – asymmetric striding in the field to stay in one place on the slow current, play speed 1x; 10-25 s. – asymmetric striding in the field, slowed down 4x; 26-34 s. – asymmetric striding in the field, view from above, slowed down 4x.

**Movie S6. *G. gigas* turning in the field**

1-4 s. – turning in the field, slowed down 4x.

**Movie S7. *C. costalis* striding in the field**

1-5 s. – asymmetric striding in the field, play speed 1x; 6-15 s. – asymmetric striding in the field, slowed down 3x.
